# Convergence stability, uninvadability, and mutual invadability for function-valued traits and life-history schedules on Hilbert spaces

**DOI:** 10.64898/2026.11.21.768256

**Authors:** Ludovic Maisonneuve, Piret Avila, Laurent Lehmann

**Affiliations:** Department of Social and Behavioral Sciences, Toulouse School of Economics, Université Toulouse Capitole, Toulouse, France; Faculty of Biological and Environmental Sciences, University of Helsinki, 00014 Helsinki, Finland; Department of Ecology and Evolution, University of Lausanne, 1015 Lausanne, Switzerland

**Keywords:** adaptive dynamics, function-valued traits, life-history traits, optimal control, convergence stability, evolutionary branching

## Abstract

The expression of many biological traits depends on environmental conditions, physiological state, or age. Such traits are naturally described as function-valued traits. Conditions determining whether their gradual evolution leads to a single stable phenotype or to phenotypic diversification have been suitably established for finite-dimensional traits. Yet, function-valued traits can also be infinite-dimensional, as with continuous age-structure, and their analysis remains challenging. Here, we extend invasion analysis to infinite-dimensional function-valued traits. We derive conditions for identifying candidate evolutionary endpoints, determining whether selection draws nearby populations toward them (convergence stability), and assessing whether, once reached, these endpoints resist invasion by nearby mutants (local uninvadability) or may instead allow evolutionary branching. These conditions depend on the selection gradient, the Jacobian of the selection gradient, and the Hessian of invasion fitness. For finite-dimensional traits, a negative-definite Jacobian of the selection gradient guarantees convergence stability regardless of genetic covariances, while a negative-definite Hessian of invasion fitness guarantees uninvadability. In infinite dimensions, we show that these sufficient conditions require strong rather than ordinary negative definiteness. We then focus on the particular case of life-history traits, for which investment at one age can alter physiological states and thereby affect fitness at later ages. Using optimal-control theory, we derive expressions for the Jacobian and Hessian operators that account for these indirect effects. We illustrate our theoretical results with two models of floral investment, with and without state dynamics. Individual-based simulations complement these analyses, showing that evolutionary branching can indeed occur when the theory identifies it as a possibility.

## 1 Introduction

Many phenotypic traits expressed by organisms vary systematically with age, physiological state, or external biotic and abiotic conditions. When phenotypic expression varies with one or several underlying variables, the trait is naturally represented as a function rather than as a scalar. Such function-valued traits include reaction norms with respect to environmental gradients, age-specific life-history schedules determining growth trajectories, and behavioral rules affecting how individuals respond to each other. When the argument of the function varies continuously, the corresponding trait space is infinite-dimensional, because an evolutionary change can in principle modify phenotypic expression at every value of the argument. Yet the theory used to classify long-term evolutionary outcomes, and in particular to predict when gradual trait evolution leads to stable endpoints or to diversification, has been developed mainly for traits described by one or a finite number of values (e.g., Eshel, 1983; Taylor, 1989; Lessard, 1990; Christiansen, 1991; Geritz et al., 1998; Metz et al., 1996; Eshel et al., 1997; Leimar, 2009; Sasaki and Dieckmann, 2011; Débarre et al., 2014; Geritz et al., 2016; Mullon and Lehmann, 2019).

This theory describes gradual evolutionary change under mutations of small phenotypic effect through the vector of derivatives of a relevant fitness measure—the selection gradient. A central concept is that of a singular point ***u***^*\**^ *∈* ℝ^*d*^, a trait value in the interior of the trait space at which the selection gradient vanishes, so that directional selection is absent. A singular point is thus a candidate endpoint of evolution, but whether it can be approached, and what happens once it is reached, remain to be determined. Two conceptually distinct stability properties must then be considered.

The first is convergence stability, which determines whether successive substitutions by mutants of small phenotypic effect tend to move a nearby resident population toward ***u***^*\**^ (convergence stable) or whether ***u***^*\**^ is unstable. Locally, this property is determined by how the selection gradient changes around the singular point, represented by its Jacobian matrix together with the genetic variance–covariance matrix, which describes how heritable variation is distributed across phenotypic directions (Leimar, 2009; Débarre et al., 2014; Geritz et al., 2016). When the symmetric part of the Jacobian matrix is negative definite, ***u***^*\**^ is a local attractor of the evolutionary dynamics for every positive definite genetic variance–covariance matrix (i.e., ***u***^*\**^ is locally exponentially stable), whereas if it is not negative semidefinite, there exists a covariance matrix for which ***u***^*\**^ is convergence unstable (Leimar, 2009).

Even if evolution approaches a singular point, a second question remains: can nearby mutants invade once it is reached? This property, which we refer to as local uninvadability, depends on the Hessian matrix of the fitness measure at hand with respect to the mutant trait. If this Hessian is negative definite, ***u***^*\**^ is locally uninvadable, whereas if it is not negative semidefinite, some nearby mutants can invade (Lessard, 1990; Leimar, 2009; Geritz et al., 2016). Combining convergence stability and local uninvadability helps determine the long-term evolutionary fate of a population. A singular point that is both convergence stable and locally uninvadable represents a stable endpoint of gradual evolution, where monomorphism is maintained. In contrast, when a singular point is convergence stable but locally invadable, nearby mutants can potentially coexist and diverge, making it a candidate evolutionary branching point (Geritz et al., 1998, 2016). When a singular point is convergence unstable, evolutionary trajectories may instead move away from it along some phenotypic directions. Thus, classifying singular points requires considering both their convergence stability and their local uninvadability, as well as the genetic variance–covariance matrix when traits are multidimensional.

How do these concepts extend to the realm of infinite-dimensional traits? The earliest evolutionary analyses of infinite-dimensional function-valued traits arose in quantitative genetics, where morphological shapes and growth trajectories were explicitly treated as infinite-dimensional characters (e.g., Kirkpatrick and Heckman, 1989; Gomulkiewicz and Kirkpatrick, 1992; Gomulkiewicz and Beder, 1996; Beder and Gomulkiewicz, 1998). Function-valued traits are also central to life-history theory, where they specify how an organism allocates resources to growth, maintenance, or reproduction over its lifespan. In this setting, approaches based on optimal control theory, such as Pontryagin’s maximum principle and dynamic programming, have long been used to identify singular functions ***u***^*\**^ *∈ U*_ad_, where *U*_ad_ is a set of admissible trait functions, under a wide range of different life-history evolution scenarios (e.g., León, 1976; Iwasa and Roughgarden, 1984; Perrin, 1992; Perrin and Sibly, 1993; Day and Taylor, 2000; Houston and McNamara, 1999; Mangel et al., 1988; Ewald et al., 2007; Nakamura and Ohtsuki, 2016; Avila et al., 2021).

The stability properties of infinite-dimensional singular functions have also been considered. Dieckmann et al. (2006) introduced function-valued counterparts of the Jacobian and Hessian and stated conditions for convergence stability and local uninvadability in terms of them, without providing proofs, whereas Parvinen et al. (2006, 2013) characterized singular functions for traits with direct effects on fitness and for traits acting through state dynamics, respectively. What remains to be established, however, is a justification of these stability criteria in function spaces. In infinite dimensions, convergence stability and uninvadability must be assessed over infinitely many phenotypic directions, where conditions based on eigenvalues and definiteness do not carry over directly from finite dimensions. This requires establishing conditions on Jacobian and Hessian operators that imply convergence stability and local uninvadability in function space, together with conditions under which convergence toward a singular function is accompanied by mutual invadability and thus creates the conditions for evolutionary diversification.

Here, we address this gap by deriving conditions that classify singular functions arising in the adaptive dynamics of function-valued traits. We first formulate evolutionary invasion analysis in a function space and characterize candidate evolutionary endpoints through the functional selection gradient. We then show that strong (i.e., uniform) negative definiteness of the Jacobian operator of this gradient ensures convergence stability for any positive definite genetic variance–covariance operator, and that strong negative definiteness of the Hessian operator of mutant invasion fitness ensures local uninvadability, together with converse conditions for instability and invadability. By combining these operators, we obtain an infinite-dimensional classification of singular functions and establish conditions under which mutually invadable functions occur arbitrarily close to a convergence-stable singular function, thereby identifying candidate evolutionary branching points. We then apply these results to analyze selection on life-history traits whose effects on fitness are mediated by age-dependent state dynamics. Building on the standard optimal-control representation of the selection gradient as the derivative of the Hamiltonian function, we derive the corresponding Jacobian and Hessian operators while accounting for changes in the state and costate trajectories. We illustrate our results with two models for the evolution of a seasonal floral investment schedule, with and without state dynamics, and use individual-based simulations to examine the diversification dynamics predicted by the local analysis.

## 2 Evolutionary invasion analysis for function-valued traits

### 2.1 Model assumptions and notation

We consider a population of infinite size of clonally reproducing individuals. Each individual expresses a heritable trait that is a function, ***u*** : *A → Z*. The argument 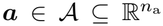 is described by *n*_a_ variables, such as age, physiological state, or environmental conditions. The codomain 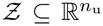 is the set of *n*_u_ phenotypic values the individual can express at any such point. This *function-valued trait* thus assigns to every argument ***a*** *∈ A* a vector ***u***(***a***) *∈ Z*. Throughout, bold symbols denote vectors, matrices, and vector- or matrix-valued functions or kernels. For a vector ***y*** *∈* ℝ^*m*^ depending on a vector ***z*** *∈* ℝ^*n*^, we define *∂****y****/∂****z*** as the *n × m* matrix whose entry (*i, j*) is *∂y*_*j*_*/∂z*_*i*_. Thus, the derivative of a scalar with respect to a vector is a column vector. The argument may vary continuously over a domain 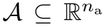, or take values in a discrete subset of 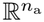; we treat both cases together. Biologically, the argument ***a*** may represent an environmental variable, such as temperature. In this case, ***u*** is a reaction norm specifying the phenotype expressed across environmental conditions. Alternatively, ***a*** may represent an individual state, such as age or body size, in which case ***u***(***a***) specifies how the expressed phenotype varies as the individual moves through different ages or states.

We assume that individual traits belong to the Hilbert space

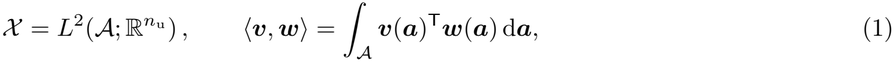

with associated norm 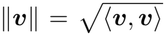, where 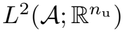 denotes the space of square-integrable functions from *A* to 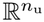 . The superscript T denotes transposition, so ***v***(***a***)^⊤^***w***(***a***) is the inner product on 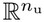. When the argument ***a*** *∈ A* varies continuously, the integral is taken with respect to the Lebesgue measure. When the argument ***a*** takes discrete values, we interpret it as

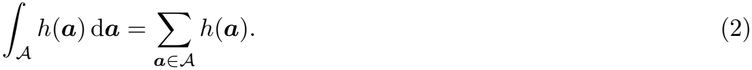

We use the integral notation for both cases throughout.

Not every function in *X*, however, necessarily represents a biologically feasible phenotype. Biological constraints may, for example, restrict the values of ***u***(***a***) to an admissible set. We therefore define *U*_ad_ *⊆ X* as the set of biologically admissible trait functions. When the argument varies continuously, we further require functions in *U*_ad_ to be bounded and piecewise continuous, meaning that *A* can be partitioned into finitely many regions on each of which the function is continuous.

Our aim is to provide a recipe to both identify singular functions ***u***^*\**^ *∈ U*_ad_ and provide a local description of their convergence stability and uninvadability, i.e., to extend the study of singular points (Geritz et al., 1998) to the infinite-dimensional case. For this aim, it is sufficient to use evolutionary invasion analysis, which characterizes the evolutionary dynamics from the invasion success of a rare mutant in a monomorphic resident population (e.g., Eshel, 1983; Taylor, 1989; Christiansen, 1991; Geritz et al., 1998; Metz et al., 1996; Eshel et al., 1997; Avila and Mullon, 2023; Van Cleve, 2023). This success is measured by the mutant’s invasion fitness *ρ*(***u***_m_, ***u***_r_), here defined as the factor deciding whether a mutant lineage initiated by a single individual bearing ***u***_m_ in a resident population bearing ***u***_r_ has a positive probability of invading the population when *ρ*(***u***_m_, ***u***_r_) *>* 1 and dying out with certainty when *ρ*(***u***_m_, ***u***_r_) *≤* 1, under the assumption that the multiplicative growth process of the mutant lineage in the resident population can be envisioned as a branching process (Crump and Mode, 1968; Mode, 1971; Tanny, 1981; Haccou et al., 2005). When the mutant invades, *ρ*(***u***_m_, ***u***_r_) is the asymptotic per-capita growth factor of the mutant lineage, i.e., *ρ*(***u***_m_, ***u***_r_) is the factor by which the mutant population is multiplied per generation (discrete time, while log(*ρ*(***u***_m_, ***u***_r_)) gives the growth rate in continuous time). We assume that invasion fitness is defined on an open subset of *X × X* containing all pairs of admissible traits and three times continuously Fréchet differentiable. Biologically, this regularity assumption means that sufficiently small changes in the mutant or resident trait produce smooth changes in invasion fitness, without jumps or sharp thresholds.

To carry out our analysis, we make two standard invasion analysis assumptions. First, mutations are rare, so the resident population reaches a locally asymptotically stable demographic equilibrium before a mutant arises. We can then study the mutant in the resident environment at this equilibrium, which implies *ρ*(***u***_r_, ***u***_r_) = 1. Second, mutations have weak phenotypic effects, so that for each resident ***u***_r_ *∈ U*_ad_ a mutant trait can be written as

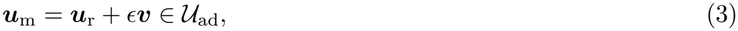

where the small parameter *ϵ* measures the magnitude of the mutational change and ***v*** *∈ X* describes the direction of the phenotypic change. Under these assumptions, we provide three criteria. First (i), we give the criterion that characterizes *candidate endpoints* of evolution ***u***^*\**^. Such candidate endpoints are of two kinds: either they lie in the interior of the phenotypic space or at the boundary. Second (ii), we give a sufficient condition for a singular function to be *convergence stable* for any genetic variance–covariance structure, ensuring that nearby evolutionary trajectories tend toward ***u***^*\**^ regardless of how genetic variation is distributed across the trait. Third (iii), we ask whether a convergence-stable ***u***^*\**^ is locally *uninvadable*. If no sufficiently close mutant can invade, the population stays monomorphic and ***u***^*\**^ is an evolutionary endpoint. If some can, mutually invadable traits may exist around ***u***^*\**^, potentially creating the conditions for further trait diversification, i.e., evolutionary branching (Geritz et al., 1998). Combining criteria (ii) and (iii), we give a classification of singular functions.

### 2.2 Directional selection and candidate endpoints

In order to characterize candidate endpoints of evolution, let us consider a nearby mutant function (eq. 3). First-order Taylor expanding mutant invasion fitness *ρ*(***u***_m_, ***u***_r_) around *ϵ* = 0, and then using eq. (A.15) of Engel and Dreizler (2013) to express the first-order term, gives

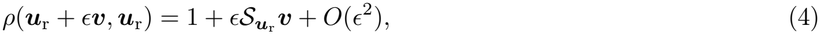

where 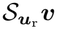 is the first-order effect of a phenotypic change in direction ***v*** on mutant invasion fitness. Because *ρ* is Fréchet differentiable, 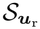 is a bounded linear functional on *X* . It can therefore be represented by a unique selection-gradient function ***s***(***u***_r_) = *{****s***(***u***_r_, ***a***)*}*_***a****∈A*_ *∈ X* (Engel and Dreizler, 2013, *eq. (A.15)), such that*

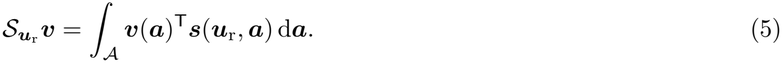

Thus, ***s***(***u***_r_, ***a***) describes how selection acts on the phenotype expressed at argument value ***a***, and 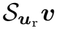 can be thought of as the directional derivative of mutant invasion fitness in direction ***v***, and 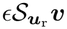 as a selection coefficient.

To calculate the selection gradient at a particular argument value ***a***, we use a perturbation concentrated around ***a***. Formally, this gives

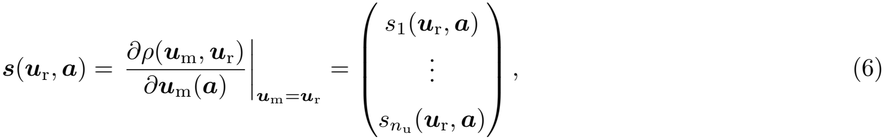

where

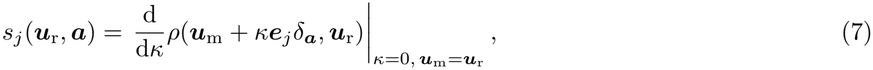

where ***e***_*j*_ is the *j*th unit vector. The derivative in eq. (6) is an ordinary partial derivative when *A* is discrete and a functional derivative when the argument varies continuously, and we use the same notation for both throughout. When *A* is discrete, *δ*_***a***_ is an indicator function, equal to one when ***a′*** = ***a*** and zero otherwise. When the argument ***a*** varies continuously, *δ*_***a***_ denotes the Dirac delta centered at ***a***, defined by *δ*_***a***_(***a***^***′***^) = *δ*(***a′****−* ***a***). Since the Dirac delta is not square-integrable, ***u***_m_ + *κ****e***_*j*_*δ*_***a***_ does not belong to *X* . Equation (7) is therefore understood as the limit obtained by replacing *δ*_***a***_ with regular functions increasingly concentrated around ***a***, taking the derivative with respect to *κ* before the limit (Engel and Dreizler, 2013, eq. (A.28)).

We refer to the preferential spread of phenotypic changes with positive first-order effects on invasion fitness as *directional selection*. Locally, *s*_*j*_(***u***_r_, ***a***) *>* 0 means that directional selection favors an increase in component *j* at ***a***, whereas *s*_*j*_(***u***_r_, ***a***) *<* 0 means that it favors a decrease. More generally, 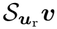 sums the first-order fitness effects of the changes specified by ***v*** across all phenotypic components and argument values, and thus describes the net effect of directional selection on the mutant trait value.

Whether directional selection can promote a change in the trait depends on the position of the resident within *U*_ad_. We call a trait function *interior* if sufficiently small changes in every direction remain admissible, and a *boundary* function otherwise. To ensure that this distinction reflects biological constraints rather than the regularity conditions imposed on trait functions, we consider only directions within the ambient function space. When the argument ***a*** varies continuously, these directions are restricted to bounded piecewise-continuous functions.

A function ***u***^*\**^ is a candidate endpoint of evolution if directional selection does not favor any admissible change away from it, that is, if

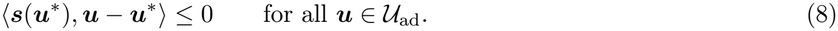

When ***u***^*\**^ is interior, sufficiently small changes along every direction considered above are admissible. The inequality must therefore hold for both a change and its opposite, which requires the selection gradient to vanish

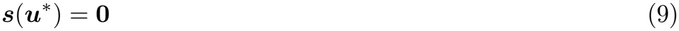

or equivalently *s*_*j*_(***u***^*\**^, ***a***) = 0 for all *j* and ***a*** *∈ A*, so that for a continuous argument, this equality holds almost everywhere in *A*. We call such an interior function a *singular function*.

**Result 1** (Candidate endpoints). *If* ***u***_r_ *is not a candidate endpoint (i.e*., *it does not satisfy the first-order condition in eq*. (8) *with* ***u***^*\**^ = ***u***_r_*), then there exists an admissible mutant function arbitrarily close to* ***u***_r_ *whose invasion fitness exceeds one because of the first-order effect of directional selection. The resident function therefore cannot be an evolutionary endpoint*.

The proofs of this and all subsequent results are given in the Appendix.

### 2.3 Convergence stability

The first-order condition (8) allows us to locate candidate evolutionary endpoints, but it does not determine whether evolution approaches any such candidate from within a neighborhood. To address this question, we combine the quantitative-genetic description for function-valued traits (Kirkpatrick and Heckman, 1989; Beder and Gomulkiewicz, 1998) with quantitative-genetic descriptions linking changes in mean trait values to invasion fitness under weak selection in the sense of small population variances (Lande, 1982; Sasaki and Dieckmann, 2011; Débarre et al., 2014; Lion, 2018; Mullon and Lehmann, 2019). When *A* is a singleton, ***ū*** _r_ is finite-dimensional, and the latter results predict that the resident mean trait ***ū***_*t*_ changes over evolutionary time *t* in proportion to *G****s***(***ū*** _*t*_), where *G* is a variance–covariance matrix. An analogous evolutionary equation has been derived for a particular model where ***u***_r_ is infinite-dimensional, where *A* can be any subset of 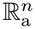 (Beder and Gomulkiewicz, 19*98). But establishing this relationship with invasion fitness in the general case remains an open problem. We therefore assume this relationship can be extended to the general case, i.e. applying to invasion fitness and A* can be any subset of 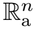, so that the mean values for the resident function ***ū*** _*t*_ evolves according to

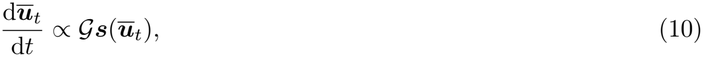

where *G* is the genetic variance–covariance operator, the function-valued analogue of the genetic variance–covariance matrix. It describes how genetic variation in trait expression at one argument value is correlated with variation at other argument values. We define this operator by

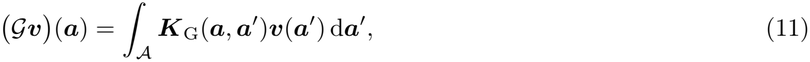

where ***K***_G_(***a, a***^***′***^) is the genetic covariance between trait values expressed at ***a*** and ***a***^***′***^. We assume that this covariance structure remains fixed throughout evolution and each component is small. Because ***K***_G_ is a covariance kernel, *G* is self-adjoint and positive semidefinite (Kirkpatrick and Heckman, 1989, Appendix A1).

We further assume that *G* is trace-class (or nuclear), as expected for the covariance operator of a random function with finite second moment (Bosq, 2000, Theorem 1.7, p. 37). Since *G* is positive and self-adjoint, this is equivalent to requiring Tr(*G*) *< ∞*, which corresponds to finite total genetic variance (where Tr is the trace). When the argument space *A* contains a finite number of discrete elements, *G* reduces to a finite variance–covariance matrix and is therefore automatically trace-class.

We now restrict attention to a singular function ***u***^*\**^. To determine whether evolution approaches such a ***u***^*\**^, we approximate the selection gradient for a resident function ***u***_r_ close to ***u***^*\**^ as

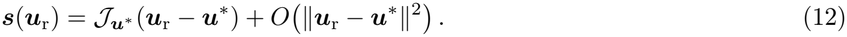

The action of the Jacobian operator 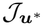 on the displacement ***u***_r_ *−* ***u***^*\**^, evaluated at ***a*** *∈ A*, can be written as

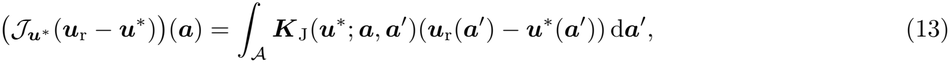

where, for each pair of arguments (***a, a***^***′***^),

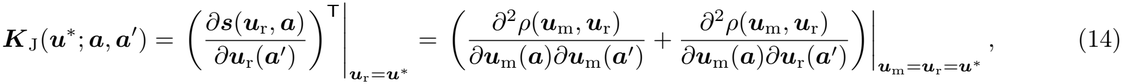

is a matrix-valued kernel in 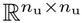 whose entry (*j, l*) measures how the displacement of resident phenotypic component *l* from its singular value at ***a′*** contributes to the selection gradient on phenotypic component *j* at ***a***. When *A* is discrete and finite, the operator 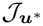 reduces to the Jacobian matrix of the selection gradient of multivariate scalar trait evolution (Leimar, 2009; Geritz et al., 2016).

Substituting eq. (12) into eq. (10) gives

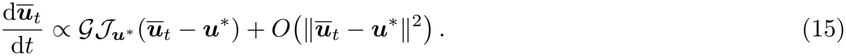

We assume that the higher-order terms *O ∥****ū***_*t*_ *−* ***u***^*\**^*∥*^2^ do not alter the local convergence properties of the evolutionary dynamics. Neglecting these terms gives the linearized evolutionary dynamics. We say that an interior singular function ***u***^*\**^ is convergence stable if the linearized evolutionary dynamics are strongly stable (Eisner, 2010, Definition 3.1, p. 100), meaning that

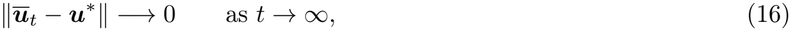

for any sufficiently nearby initial resident function. For finite-dimensional traits, strong stability of the linearized dynamics is equivalent to their exponential stability (Engel and Nagel, 2000, Proposition 3.12). In infinite-dimensional spaces, however, strong stability of the linearized dynamics does not necessarily imply their exponential stability (Engel and Nagel, 2000, Example 1.3(i)), yet it implies that ***u***^*\**^ is a local attractor of gradual evolution. Convergence stability also depends on both the Jacobian operator 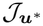 and the genetic variance–covariance operator *G*, as the following result shows.

**Result 2** (Convergence stability). *If the Jacobian operator* 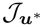 *is strongly negative definite, then* ***u***^*\**^ *is convergence stable for any positive definite genetic variance–covariance operator G. Strong negative definiteness means that there exists a constant c*_J_ *>* 0 *such that*

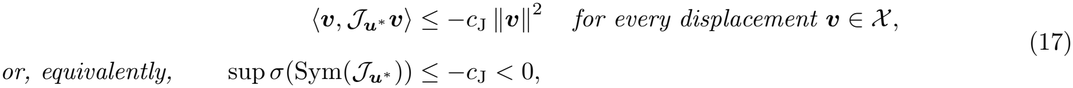

*where* 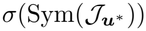 *denotes the spectrum of the symmetric part of the Jacobian operator. Conversely, if* 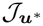 *is not negative semidefinite, there exists a positive definite genetic variance–covariance operator for which* ***u***^*\**^ *is unstable. When* 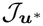 *is negative semidefinite but not strongly negative definite, no general conclusion follows (see Appendix A.2 for a reminder of the definitions of definiteness)*.

For finite-dimensional traits, the spectrum 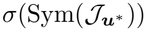 consists of the eigenvalues of the symmetric part of the Jacobian matrix, in which case ***u***^*\**^ is convergence stable if the leading eigenvalue of 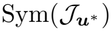 is negative. Result 2 without its last statement then reduces to that of Leimar (2009, p. 198, where convergence stability is called strong convergence stability). In infinite dimensions, 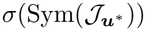 may also contain values that are not eigenvalues. The symbol sup in (17) denotes the largest value in this spectrum, or its least upper bound if no largest value exists. Thus, the condition in eq. (17) requires all functional modes to have a negative restoring effect, whose strength is uniformly bounded below by *c*_J_.

### 2.4 Uninvadability and polymorphism

Convergence stability determines whether evolutionary trajectories starting sufficiently close to ***u***^*\**^ are drawn toward it. We now ask whether a population that has reached ***u***^*\**^ will remain there so that ***u***^*\**^ can be envisioned as a final local endpoint of gradual evolution. This will depend on whether it can be invaded by nearby mutant functions. Because directional selection vanishes for a singular function 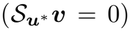, the second-order Taylor expansion of invasion fitness for a mutant function ***u***^*\**^ + *ϵ****v*** satisfies

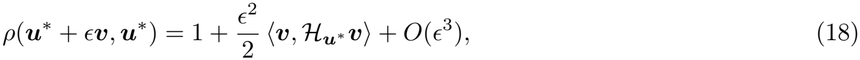

where 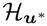 is the Hessian operator at ***u***^*\**^ whose action and matrix-valued kernel are

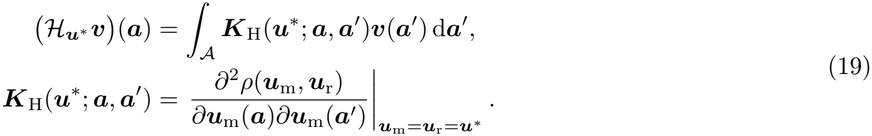

Biologically, each element [***K***_H_(***u***^*\**^; ***a, a***^*′*^)]_*j.l*_ describes the synergistic fitness effect of jointly changing phenotypic component *j* at argument value ***a*** and component *l* at argument value ***a***^***′***^. When *A* is discrete and finite, the operator 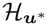 reduces to the Hessian matrix of invasion fitness of multivariate scalar trait evolution (e.g., Lessard, 1990; Leimar, 2009; Geritz et al., 2016).

An interior singular function ***u***^*\**^ will be said to be locally uninvadable if it is a strict local maximum of mutant invasion fitness, such that any sufficiently small admissible deviation from ***u***^*\**^ has invasion fitness below one.

**Result 3** (Uninvadability). *A sufficient condition for a singular function* ***u***^*\**^ *to be locally uninvadable is that* 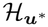 *be strongly negative definite. That is, there exists a constant c*_H_ *>* 0 *such that*

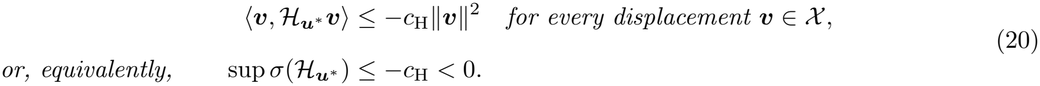

*Mutant invasion fitness then decreases in every functional direction, making* ***u***^*\**^ *a strict local fitness maximum. Conversely, if* 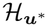 *is not negative semidefinite, then* ***u***^*\**^ *is locally invadable. If* 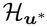 *is negative semidefinite but not strongly negative definite, the second-order analysis alone is inconclusive*.

Note that the Hessian operator is self-adjoint because it represents the symmetric second derivative of invasion fitness with respect to two mutant perturbations. Unlike the Jacobian, it therefore does not need to be replaced by its symmetric part before its spectrum is considered. For finite-dimensional traits, ***u***^*\**^ is locally uninvadable if the leading eigenvalue of 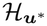 is negative and invadable if at least one eigenvalue of 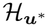 is positive.

When a singular function is both convergence stable and locally uninvadable, nearby evolutionary trajectories approach it, and no sufficiently similar mutant can invade once it is reached. It is therefore a locally evolutionarily stable endpoint of the gradual evolution process. But what happens when a singular function is convergence stable but does not satisfy the sufficient condition for uninvadability?

**Result 4** (Mutual invadability). *If an interior singular function* ***u***^*\**^ *satisfies the condition in eq*. (17) *but does not satisfy the uninvadability condition in eq*. (20), *then there exist two arbitrarily close traits on opposite sides of* ***u***^*\**^ *that are mutually invadable*.

For finite multidimensional traits, Geritz et al. (2016) characterized the conditions under which two traits near a singular trait can mutually invade each other, depending on their positions relative to that trait. In particular, considering traits symmetrically positioned around the singular trait, their analysis implies that a negative definite Jacobian matrix and a Hessian matrix that is not negative definite are sufficient for mutual invadability. Result 4 extends this sufficient condition to function-valued traits, using the corresponding criteria based on strong negative definiteness. Mutual invadability does not guarantee, however, that such mutations will actually arise, establish, and generate the emergence of two coexisting phenotypic lineages that subsequently diverge. Whether this evolutionary branching will actually occur can then be determined with individual-based simulations with recurrent mutations (and see also Geritz et al., 2016).

## 2.5 Classification of singular functions

The Jacobian 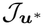 and Hessian 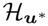 operators characterize two distinct properties of a singular function. The Jacobian determines whether selection acting on nearby resident functions brings them toward ***u***^*\**^, whereas the Hessian determines whether nearby mutants can invade a population expressing ***u***^*\**^. Combining these properties yields the following five cases.

1. When both 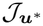 and 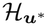 are strongly negative definite, nearby resident functions are brought back toward ***u***^*\**^ for any positive definite genetic variance–covariance operator *G*, and no sufficiently similar mutant can invade once it is reached. The singular function is therefore a local attracting endpoint of gradual evolution, and is what corresponds to the biological intuition of an evolutionary stable trait value (e.g., Maynard Smith, 1982), also called a *continuously stable trait* in the finite-dimensional case (Eshel, 1983; Lessard, 1990).
2. When 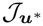is strongly negative definite and 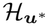 is negative semidefinite but not strongly negative definite, nearby resident functions still approach ***u***^*\**^ for any positive definite genetic variance–covariance operator *G*, but the second-order analysis does not determine whether ***u***^*\**^ itself is invadable. Nevertheless, Result 4 guarantees the existence of mutually invadable trait functions arbitrarily close to it, so nearby polymorphism is possible. The singular function is therefore a candidate *evolutionary branching point*. It is only a candidate because mutual invadability does not by itself guarantee the establishment, coexistence, and subsequent divergence of two phenotypic lineages.
3. When 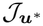 is strongly negative definite but 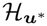 is not negative semidefinite, nearby resident functions approach ***u***^*\**^ for any positive definite genetic variance–covariance operator *G*, while Result 3 implies that ***u***^*\**^ can be invaded and Result 4 guarantees mutually invadable trait functions arbitrarily close to it. The singular function is therefore also a candidate *evolutionary branching point*.
4. When 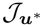 is negative semidefinite but not strongly negative definite, the present criteria do not establish whether ***u***^*\**^ can be convergence stable or unstable, irrespective of the properties of 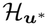 .
5. Finally, if 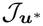 is not negative semidefinite, there exists at least one positive definite genetic variance–covariance operator for which ***u***^*\**^ is convergence unstable. This does not imply instability under every genetic variance– covariance operator.

Although the Jacobian and Hessian characterize different evolutionary properties, they are mathematically related. As shown in Appendix A.4 (see eq. (A.118)),

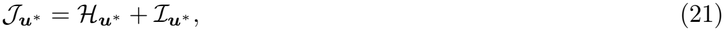

where the interaction operator 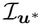 has matrix-valued kernel

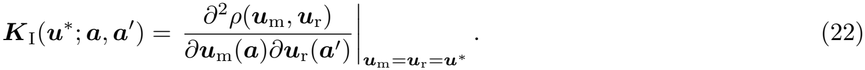

This operator measures how the marginal fitness effect of changing the mutant trait value at ***a*** depends on the trait value expressed by the resident at ***a***^***′***^, and thus captures trait-dependent selection. In the absence of trait-dependent interactions between residents and mutants, 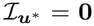, so 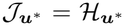. Consequently, a singular function satisfying the strong convergence stability condition is also locally uninvadable, ruling out evolutionary branching. For scalar traits, evolutionary branching requires negative trait-dependent selection, meaning that increasing the resident trait reduces the selective advantage of increasing the mutant trait (Geritz et al., 1997; Kisdi, 2015; Lehmann and Mullon, 2026). Identifying the properties of 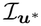 necessary for evolutionary branching in multidimensional and function-valued traits remains an open question.

## 2.6 Example: evolution of a floral investment schedule

### 2.6.1 Biological assumptions

To illustrate how the preceding criteria apply to a biological function-valued trait, we consider the evolution of flowering schedules, which have been used previously to illustrate the adaptive dynamics of function-valued traits and evolutionary branching (Dieckmann et al., 2006). Here, we formulate an alternate version of this model that replaces the fixed total flower production assumed by Dieckmann et al. (2006) with explicit reproductive costs of floral investment, making the underlying biological trade-off more transparent. In this version, a floral investment schedule *u* specifies an individual’s investment in floral production *u*(*a*) *∈* [0, *∞*) at each time *a* during a flowering season of fixed length *A* = [0, 1] (because *u*(*a*) is scalar in this example, we omit the bold notation used above for vector-valued traits).

Individuals are assumed to be haploid and reproduce asexually. They express their flowering schedule and produce clonal seeds following pollination throughout the season, but release all offspring, including vegetative propagules, simultaneously at its end, when they die, so generations do not overlap. Offspring then compete for a fixed number of establishment sites, and this competition regulates the population to a constant size. Population renewal is therefore assumed to follow a Wright–Fisher process (Crow and Kimura, 1970; Ewens, 2004).

Vegetative propagules are assumed to be produced in a baseline quantity *f*_0_ *≥* 1, which gives the expected number of offspring produced through vegetative propagation. Clonal seeds are produced through pollen-dependent apomixis (pseudogamy), in which pollen is required for endosperm development but does not fertilize the embryo. Investment *u*(*a*) in floral reproduction generates such seeds, e.g., by maintaining stigmatic receptivity, supporting pollen-tube growth, and provisioning developing ovules, but it also carries maintenance costs and changes how plants interact through pollinators. We assume that the total net quantity of offspring produced through pollinated seeds by a mutant is 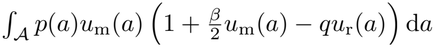. Here *p*(*a*)*u*_m_(*a*) is the baseline quantity of pollen collected by the mutant: *p*(*a*) is a continuous function describing pollinator availability, while *u*_m_(*a*) captures how effectively the plant attracts pollinators (e.g., through flower display). The parameter *β ≥* 0 captures increasing returns within the mutant’s own reproductive investment. Greater investment may improve the efficiency with which received pollen is converted into mature seeds, for example by increasing stigmatic receptivity, supporting pollen-tube growth, or provisioning developing ovules. Resident investment reduces the mutant’s return at rate *q ≥* 0 because resident and mutant displays compete for pollinators. Finally, floral investment is assumed to take a toll 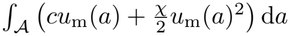 on vegetative reproduction, where *c ≥* 0 and *χ ≥* 0 control the linear and quadratic costs of reproductive investment.

Combining the different terms, the expected fecundity of a mutant expressing schedule *u*_m_ in a monomorphic resident population expressing *u*_r_ can be written as

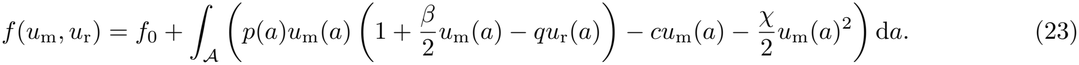

In the Wright–Fisher process, the contribution of each individual to the next generation is proportional to its expected fecundity, whereby the invasion fitness of a mutant is

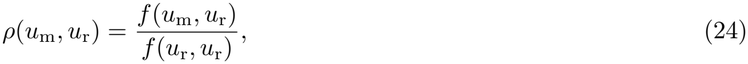

which satisfies *ρ*(*u*_r_, *u*_r_) = 1 for all *u*_r_ *∈ U*_ad_.

### 2.6.2 Directional selection and candidate endpoints

We first determine how selection affects investment in floral reproduction at each time of the season. Let *f*_r_ = *f* (*u*_r_, *u*_r_). Applying the definition of the selection gradient in eq. (6) to mutant invasion fitness eq. (24) gives

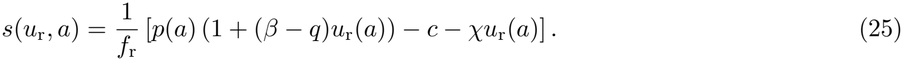

Thus, at each time in the season, selection favors an increase in floral investment if the marginal reproductive benefit of floral investment outweighs its marginal cost. Solving *s*(*u*^*\**^) = 0 in *U*_ad_ gives the singular schedule

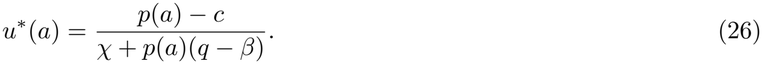

We assume that *χ* + *p*(*a*)(*q − β*) *>* 0 and *p*(*a*) *> c* throughout the flowering season. The singular schedule in eq. (26) is then strictly positive and therefore entirely interior. Because *p* is continuous, *u*^*\**^ is continuous and therefore admissible.

### 2.6.3 Convergence stability

We next ask whether selection brings nearby resident schedules toward *u*^*\**^. Differentiating the selection gradient (25) with respect to the resident schedule and evaluating it at *u*^*\**^ gives the Jacobian kernel

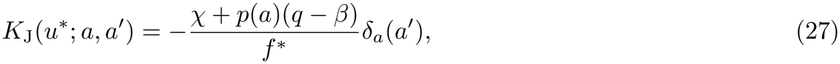

where *f* ^*\**^ = *f* (*u*^*\**^, *u*^*\**^) and *δ*_*a*_(*a*^*′*^) is the Dirac delta function centered at *a* and evaluated at *a*^*′*^. Consequently, for any functional displacement *v*,

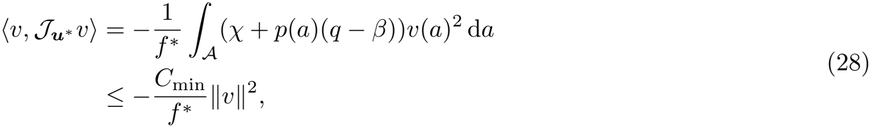

where *C*_min_ = min_*a∈A*_ (*χ* + *p*(*a*)(*q − β*)). By assumption, *χ* + *p*(*a*)(*q − β*) *>* 0 for all *a ∈ A*, so *C*_min_ *>* 0. The Jacobian is therefore strongly negative definite, implying that *u*^*\**^ determined by eq. (26) is convergence stable for any positive definite genetic variance–covariance operator.

### 2.6.4 Uninvadability and polymorphism

Finally, we determine whether nearby mutant schedules can invade once the population has reached *u*^*\**^. Differentiating the expression of invasion fitness given in eq. (24) twice with respect to mutant investment gives the Hessian kernel

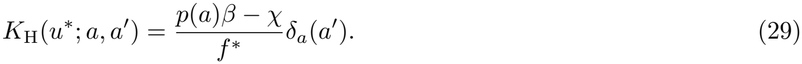

Its effect along a functional displacement *v* is therefore

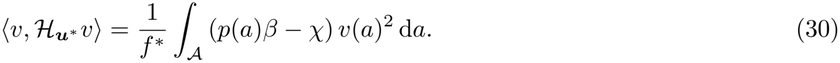

If *p*(*a*)*β − χ <* 0 throughout the season, then

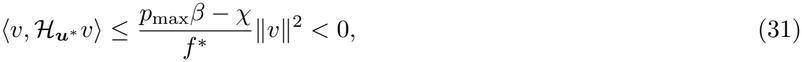

for every non-zero displacement *v*, where *p*_max_ = max_*a∈A*_ *p*(*a*). The Hessian operator is therefore strongly negative definite. Hence, *u*^*\**^ is a strict local maximum of invasion fitness and is locally uninvadable.

Conversely, if *p*(*a*)*β − χ ≥* 0 over a non-negligible portion of the flowering season, and because the Jacobian remains strongly negative definite, Result 4 guarantees the existence of nearby floral investment schedules that are mutually invadable. Pollinator competition is what allows convergence stability and invadability to occur together since

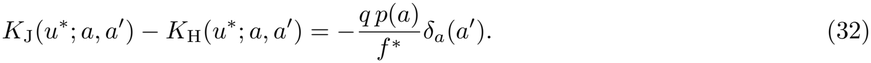

When *q* = 0, this entails 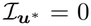 and the same local curvature determines both properties, so a singular schedule cannot be convergence stable while also being invadable. When *q >* 0, competition changes how a mutant performs against the resident without changing the mutant’s own curvature. A lineage with greater investment is favored when rare among lower-investing residents, because weak resident competition allows its larger display to benefit from the increasing returns to floral investment. Conversely, a lineage with lower investment is favored when rare among higher-investing residents, because it pays lower maintenance costs and exposes less of its display to the intense pollinator competition generated by the common strategy. Each strategy therefore loses its advantage as it becomes common, promoting their coexistence.

The local analysis identifies the selection acting near the singular schedule, but it does not establish whether recurrent mutation in a finite population actually generates and maintains distinct phenotypic lineages. We therefore complement it with individual-based simulations of the Wright–Fisher process (fig. 1, see Appendix C.1 for details). We consider two parameter sets. In the first, *p*(*a*)*β − χ <* 0 throughout the season, so the singular schedule is both convergence stable for any positive definite genetic variance–covariance operator and locally uninvadable. In the second, *p*(*a*)*β − χ >* 0 throughout the season, so the singular schedule remains convergence stable for any positive definite genetic variance–covariance operator but becomes locally invadable. In both cases, the population first approaches *u*^*\**^, as predicted by the strongly negative definite Jacobian. Under stabilizing selection, it then remains concentrated around *u*^*\**^ (fig. 1, first column of panels). Under disruptive selection, it instead splits into two groups whose floral investment schedules diverge on opposite sides of *u*^*\**^ (fig. 1, second column of panels). The simulations therefore show stabilization around a locally uninvadable singular schedule and evolutionary branching around a convergence-stable but locally invadable singular schedule, consistent with the predictions of the local analysis.

**Figure 1:**
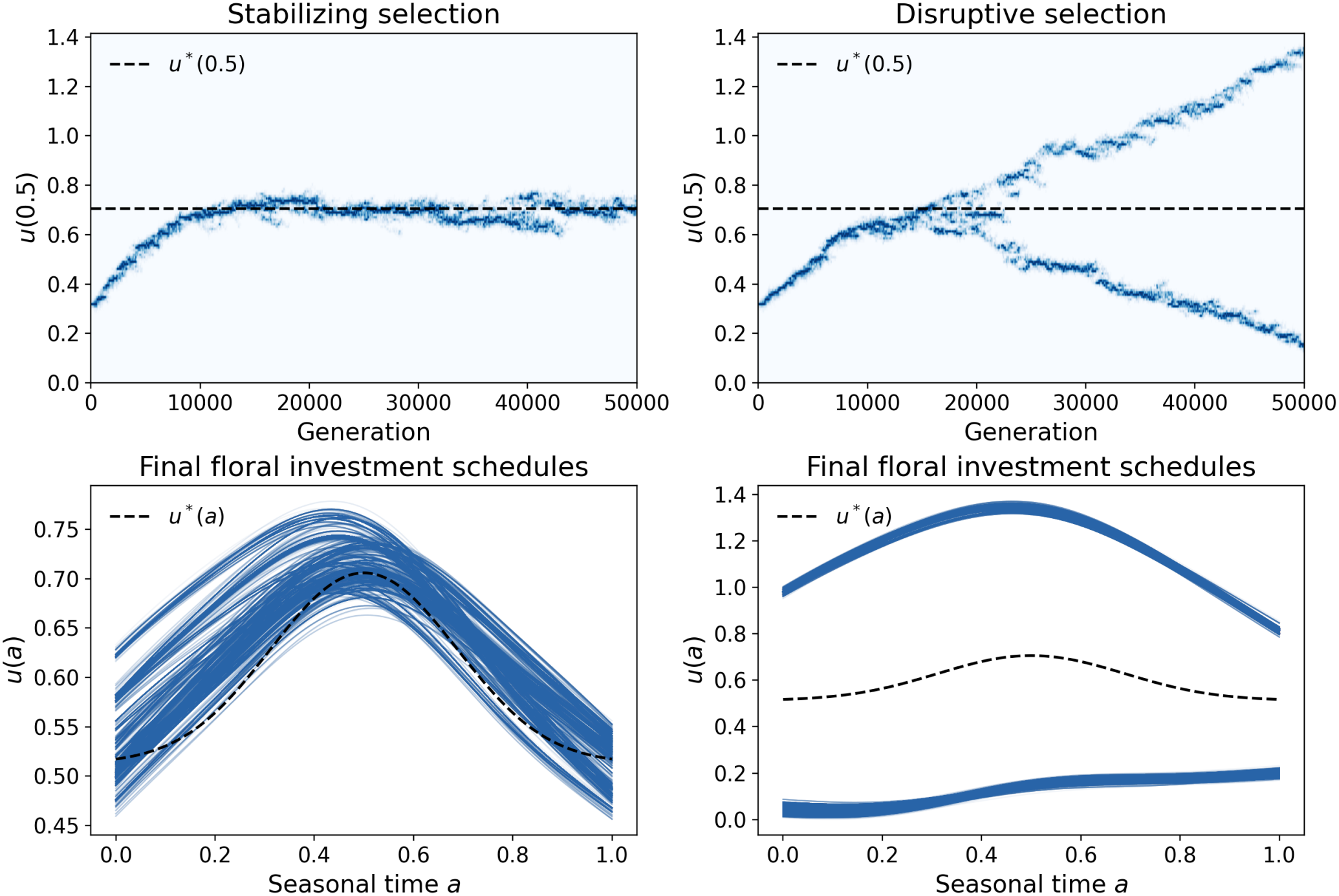
Individual-based evolution of floral investment schedules under stabilizing and disruptive selection. Top panels show the distribution of mid-season floral reproduction investment, *u*(0.5), through time, with darker blue indicating greater density. Bottom panels show the floral investment schedules expressed by all individuals in the final population. Dashed lines show the analytical singular schedule *u*^*\**^. In the stabilizing case, we use *β* = 0.35 and *q* = 0.70, whereas in the disruptive case we use *β* = 0.80 and *q* = 1.15. Common parameters were *c* = 0.80, *χ* = 1.00, *f*_0_ = 2.00, and 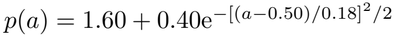. Simulation details are provided in Appendix C.1.

## 3 Analysis for life-history traits with state dynamics

The preceding analysis treats the function-valued trait as the phenotype expressed directly at each value of the argument ***a***. In many life-history problems, however, the evolving function specifies an age-specific action or allocation that changes a biological state that affects vital rates and development throughout life. Examples include allocating resources to growth, which changes body size and structures (Iwasa and Roughgarden, 1984; Kozłowski and Wiegert, 1986; Perrin, 1992; Irie and Iwasa, 2005); investing in somatic or germline maintenance, which limits the accumulation of physiological damage or deleterious mutations (Munch and Mangel, 2006; Mangel, 2008); and investing in learning, which increases acquired knowledge (Lehmann et al., 2013; Maisonneuve and Lehmann, 2026). A trait expressed at one age can therefore affect reproduction at all subsequent ages by changing the current biological state, where the effect is carried forward through life. We now extend the analysis to such life-history traits with state dynamics.

### 3.1 Model assumptions

We now consider a special case of the general framework in which the population is age-structured and the evolving trait ***u*** specifies an age-dependent control. This control affects invasion fitness through two mechanisms: directly, by changing survival or reproduction at a given age, and indirectly, by changing an individual’s biological state, which can affect survival and reproduction later in life (e.g., growth investment affecting body size, somatic or germline maintenance limiting the accumulation of physiological damage or deleterious mutations, or learning effort increasing acquired knowledge). At each age *a*, the state is a vector 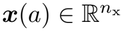 describing *n*_x_ biological attributes of the individual. Throughout this section, the argument is age, and *A* is assumed to be continuous, so that admissible control traits are piecewise continuous. An individual can shift its allocation abruptly at a finite number of developmental transitions, but not infinitely often.

Individuals thus develop, reproduce, and survive over a continuous lifespan. We denote by 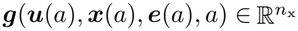the vector of rates of state change, and by *b*(***u***(*a*), ***x***(*a*), ***e***(*a*), *a*) and *µ*(***u***(*a*), ***x***(*a*), ***e***(*a*), *a*) the birth and death rates, respectively, at age *a*. Here, 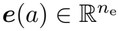 denotes *n*_e_ ecological conditions experienced by the individual at age *a*, such as population density, resource availability, or the abundance of competitors.

Mortality determines whether an individual reaches later ages. We describe this through the probability *l*(*a*) of surviving from birth to age *a*. Collecting survival and the biological state into the *extended state*

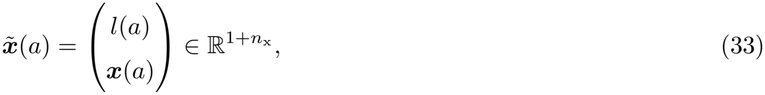

the individual develops over its lifespan according to

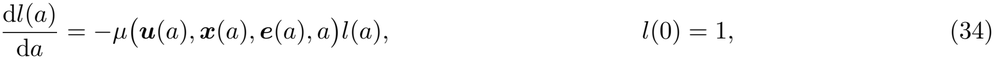

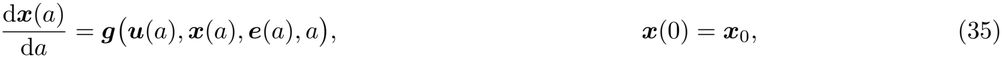

where ***x***_0_ is the vector of biological state variables at birth. The first equation describes the decline in survival probability generated by age-specific mortality, whereas the second describes how the biological state changes with age.

We write these dynamics compactly as

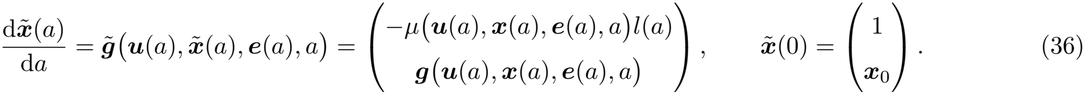

The age domain is either *A* = [0, *∞*) when lifespan is unbounded or *A* = [0, *a*_f_] when individuals cannot live beyond a maximum age *a*_f_ . In the finite-lifespan case, individuals that survive to *a*_f_ may convert their terminal state into offspring. We represent this by Φ(***x***(*a*_f_), ***e***(*a*_f_)), the expected number of offspring produced at the end of life by an individual in terminal state ***x***(*a*_f_). This term may describe, for example, a final reproductive event in a semelparous organism or the conversion of stored reserves into offspring. Its expected contribution is *l*(*a*_f_)Φ(***x***(*a*_f_), ***e***(*a*_f_)). When *A* = [0, *∞*), survival tends to zero at late ages, and reproduction is therefore entirely described by the age-specific birth rate *b*.

We now apply the analysis of section 2.1 to this age-structured setting. Consider a rare mutant expressing control trait ***u***_m_ in a monomorphic resident population expressing ***u***_r_ at its demographic equilibrium. The resident population is thus assumed to generate equilibrium ecological conditions experienced by the mutant at age *a*, which we denote by ***e***(***u***_r_, *a*), a vector of feedback variables that we do not model explicitly, as they will not be needed as such (see de Roos, 1997; Diekmann et al., 2003; Kisdi and Geritz, 2016 for explicit models). Through these conditions, ***u***_r_ can affect the mutant’s reproduction rate *b*, mortality rate *µ*, and state dynamics ***g***, and therefore its state trajectory ***x***_m_ and survival probability *l*_m_. The mutant’s basic reproductive number is

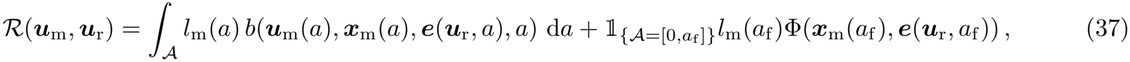

where ***x***_m_ and *l*_m_ follow the mutant state and survival dynamics in the resident population, and 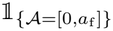 equals one when lifespan is bounded and zero otherwise. The integral gives expected reproduction throughout life, whereas the final term accounts for terminal reproduction when lifespan is bounded.

Standard results for age-dependent branching processes entail that *ρ*(***u***_m_, ***u***_r_) *≤* 1 *⇐⇒ ℛ* (***u***_m_, ***u***_r_) *≤* 1 and so the mutant lineage goes extinct if *ℛ* (***u***_m_, ***u***_r_) *≤* 1 and can invade when *ℛ* (***u***_m_, ***u***_r_) *>* 1 (Mode, 1969, 1971). Thus, *ℛ* provides an invasion fitness proxy and all the preceding results therefore remain valid when *ρ* is replaced by *ℛ*.

A change in the control expressed at age *a* perturbs the state dynamics from that age onward and therefore affects both survival and reproduction at all subsequent ages. To calculate the selection gradient ***s***, its Jacobian *J*, and the Hessian *ℋ*, we must therefore account for how these effects propagate through life. In the next section, we explain how to calculate these quantities in this setting, and all results therein are derived in Appendix B.

### 3.2 Hamiltonian representation of the evolutionary operators

#### 3.2.1 Hamiltonian function

A change in the control expressed at age *a* can affect fitness in two ways. It can directly affect reproduction or survival at that age, or perturb the extended biological state, thereby altering its dynamics over the rest of life and affecting future survival and reproduction. The Hamiltonian combines these effects through two terms: current reproduction and the change in future reproductive output generated by the biological state,

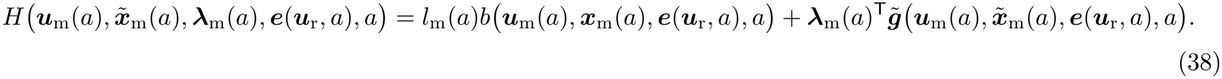

The vector

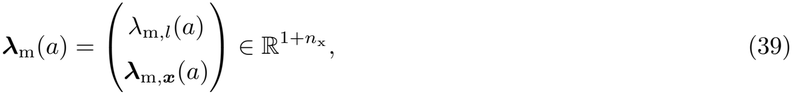

contains the mutant costate variables, whose components measure the marginal effects of the state variables 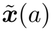 on the mutant’s remaining expected reproductive output, with *λ*_m,*l*_(*a*) being Fisher’s reproductive value. The costate vector ***λ***_m_(*a*) is defined by the backward dynamics

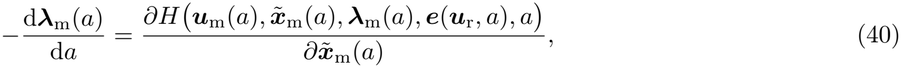

where 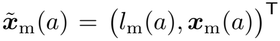 denotes the mutant extended state at age *a*, defined by 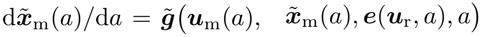, and 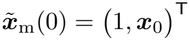. When *A* = [0, *a*_f_], its terminal condition is determined by the reproductive gain obtained from the terminal state,

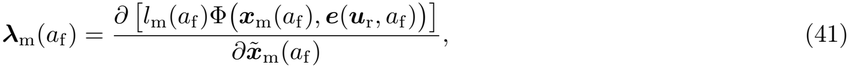

which thus provides a boundary condition for the costate dynamics (40). When *A* = [0, *∞*), no terminal state is reached and we instead impose the transversality condition

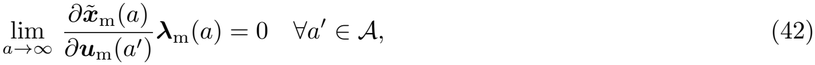

where 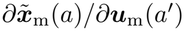 measures how a small change in the mutant control at age *a*^*′*^affects its state at age *a* (see also eq. (B.13)).

The transversality condition eq. (42) requires the effect of this change on remaining reproductive output to vanish as *a → ∞*, for every *a*^*′*^ *∈ A*. This condition helps determine the appropriate boundary conditions for the costate dynamics. In particular, because *λ*_m,*l*_(*a*) represents the expected remaining reproductive output conditional on survival to age *a*, we have *λ*_m,*l*_(0) = *ℛ* (***u***_m_, ***u***_r_).

Note that in standard optimal-control approaches to life-history evolution, the costate is typically defined along the resident trajectory, as this is sufficient to characterize singular functions (Day and Taylor, 1997, 2000; Metz et al., 2016; Avila et al., 2021). Here, we instead define the costate along the mutant trajectory, which allows us to obtain a compact expression for the Hessian operator (Result 7 below).

#### 3.2.2 Selection gradient

In terms of the above quantities, we have the following representation of the age-specific selection gradient.

**Result 5** (Directional selection for control traits). *For the basic reproductive number* (37), *the pointwise selection gradient on the control expressed at age a is*

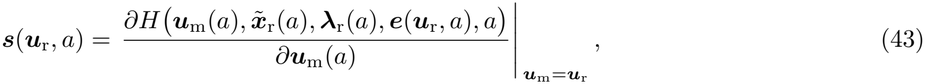

*where* ***λ***_r_(*a*) *is the resident costate, defined by eq*. (40) *with* ***u***_m_ = ***u***_r_ *and* 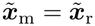, *together with the corresponding boundary condition* (41) *or* (42).

*Using the definition of the Hamiltonian, this can be written as*

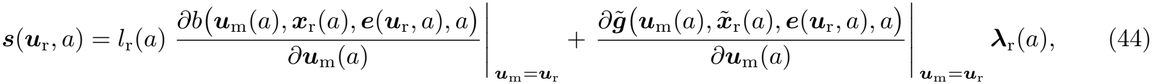

*where the matrix* 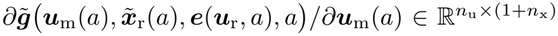 *describes how each component of the control trait affects the instantaneous dynamics of the biological state. The first term captures the effect of the control trait on current reproduction. The second term captures its effect on all subsequent reproduction through its effect on the extended state. Terminal reproduction then affects selection at earlier ages because it determines the terminal value of* ***λ***_r_, *which is propagated backward through eq*. (40).

Result 5 provides a direct procedure for computing the pointwise selection gradient: first solve the resident state dynamics, then solve the costate dynamics backward in age, and finally evaluate the derivative of the Hamiltonian with respect to the mutant control trait. This Hamiltonian representation of the selection gradient is standard in optimal-control approaches to life-history evolution (Metz et al., 2016; Avila et al., 2021). We state it here to connect these methods explicitly to our general framework for function-valued traits. Once ***s***(***u***_r_, *a*) has been obtained, Result 1 applies directly to identify candidate evolutionary endpoints.

#### 3.2.3 Jacobian operator

For an interior candidate endpoint, the Jacobian operator *J* is obtained by differentiating the selection gradient with respect to the resident control, which yields the following result.

**Result 6** (Jacobian for control traits). *At an interior singular control* ***u***^*\**^, *for the invasion fitness model* (37), *the Jacobian operator has kernel*

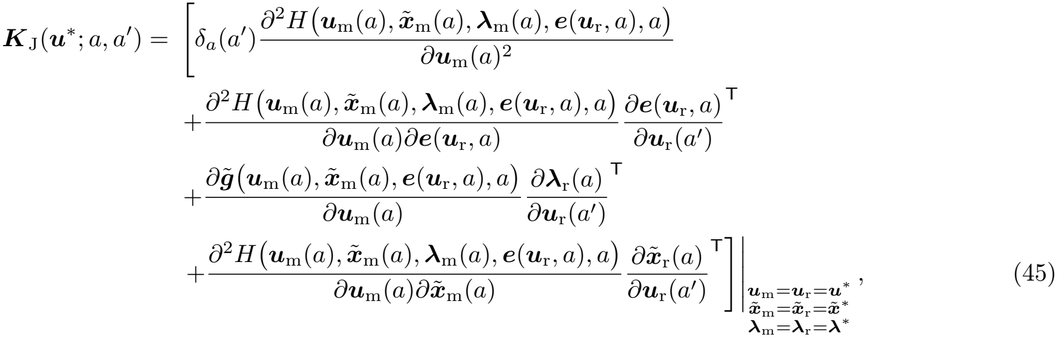

*where all derivatives are evaluated at the singular control* ***u***^*\**^ *and along the corresponding state and costate trajectories* 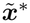 *and* ***λ***^*\**^. *The Hamiltonian derivatives with respect to the control, the state, or both have the corresponding dimensions, so that every term in* ***K***_J_(***u***^*\**^; *a, a*^*′*^) *belongs to* 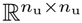.

Each matrix ***K***_J_(***u***^*\**^; *a, a*^*′*^) measures how a change in directional selection on the control at age *a* interacts with a change in resident control at age *a* (recall eq. (14)). The four terms in eq. (45) identify distinct pathways generating this interaction. The first term captures the effect due to changes expressed at the same age, through their effects on current reproduction and instantaneous state dynamics. The second term captures how a change in resident control at age *a* modifies the ecological conditions experienced at age *a* (effect 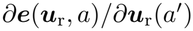), thereby altering selection on the control expressed at that age. The third term captures how a resident trait change at age *a* modifies the costate at age *a* (effect 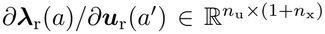), thereby changing the future reproductive value of the state changes generated by the control at that age. Unlike the state sensitivity, this can be non-zero when *a < a*^*′*^, because a change in the control at a later age can alter the future reproductive value of an earlier state change. The fourth term captures how a resident trait change at age *a* modifies the state reached at age *a* (effect 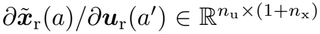), thereby altering the fitness effect of the control expressed at that age. This term vanishes when *a < a*^*′*^, because a control change cannot affect an earlier state.

Eq. (45) allows us to determine whether the singular control is convergence stable using Result 2. When explicit expressions for the resident state and costate are available as functions of the resident control, their sensitivity matrices can be obtained directly by differentiation. Otherwise, the sensitivities can be obtained by solving a system of ODEs derived in Appendix B.2. We apply this procedure later to a specific biological example in section 3.3.

#### 3.2.4 Hessian operator

We now turn to the Hessian operator, which has the following representation.

**Result 7** (Hessian for control traits). *At an interior singular control* ***u***^*\**^, *for the invasion fitness model* (37), *the Hessian operator has kernel*

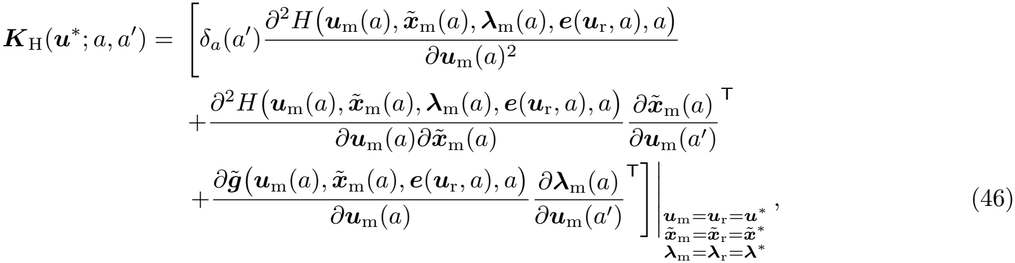

*where all derivatives are evaluated at* ***u***^*\**^ *and along the corresponding state and costate trajectories* 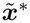 *and* ***λ***^*\**^. *The Hamiltonian derivatives with respect to the control, the state, or both have the corresponding dimensions, so that every term in* ***K***_H_(***u***^*\**^; *a, a*^*′*^) *belongs to* 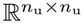.

Each matrix ***K***_H_(***u***^*\**^; *a, a*^*′*^) measures how a change in the control at age *a* interacts with a change at age *a*^*′*^ in determining mutant fitness. The first term therein captures the interaction between changes expressed at the same age, through their effects on current reproduction and instantaneous state dynamics. The second term captures how a mutant trait change at age *a*^*′*^modifies the state reached at age 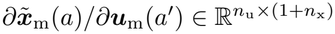, thereby altering the fitness effect of the control expressed at that age. This term vanishes when *a < a*^*′*^. The third term captures how a mutant trait change at age *a*^*′*^ modifies the costate at age *a* (effect 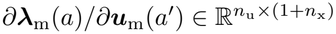, thereby changing the future reproductive value of the state changes generated by the control at that age. This sensitivity can be non-zero when *a < a*^*′*^.

Using eqs. (45)–(46), Results 3 and 4 can therefore be applied directly to determine local uninvadability or the possibility of evolutionary branching. When explicit expressions for the mutant state and costate are available as functions of the mutant control, their sensitivities can be obtained directly by differentiation. Otherwise, the sensitivities can be obtained by solving a system of ODEs derived in Appendix B.3 (in Appendix B.3.2, we provide a representation for ***K***_H_(***u***^*\**^; *a, a*^*′*^) only in terms of mutant state perturbations).

### 3.3 Example: evolution of a flower growth schedule

#### 3.3.1 Biological assumptions

To illustrate these results about how state dynamics affect the classification of singular schedules, we return to the flowering example introduced in section 2.6, under the same annual life-cycle (for which *ℛ* (*u*_m_, *u*_r_) = *ρ*(*u*_m_, *u*_r_)), but assume that floral tissue is a state variable generated by an age-dependent control. At each age, the control *u*(*a*) *∈* [0, *∞*) specifies investment in floral growth, while *x*(*a*) denotes the resulting amount of floral tissue. Investment produces floral tissue at rate *αu*(*a*), whereas existing tissue is lost at rate *νx*(*a*). The floral state therefore follows

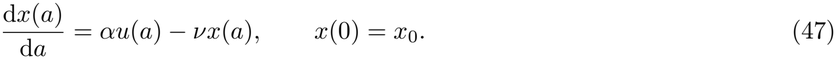

A change in investment at one age therefore alters floral display throughout the remainder of the season.

As in the preceding example, reproductive success combines benefits of attracting pollinators and producing offspring with direct costs of flower investment. Here, however, the benefits depend on the floral tissue *x*(*a*), whereas the costs depend directly on the investment in floral growth *u*(*a*). The instantaneous rate of reproductive success is therefore

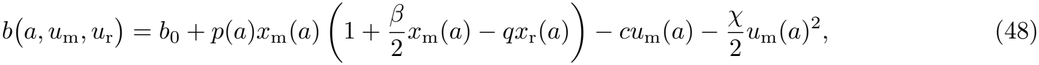

and the total mutant fecundity over the season is therefore

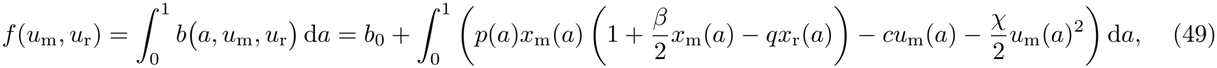

whereby relative fecundity determines invasion fitness through eq. (24).

#### 3.3.2 Directional selection and candidate endpoints

To determine the direction of selection on floral investment, we use eq. (38), which here takes the form

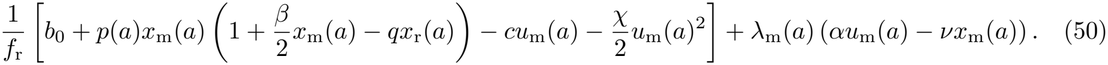

Hence, the ecological conditions generated by the resident can be written as ***e***(*u*_r_, *a*) = (*x*_r_(*a*), *f*_r_) ^⊤^.

In this example, the only state variable to consider is floral tissue *x*(*a*), because survival is fixed at *l*(*a*) = 1 throughout the flowering season. We therefore suppress the survival component from the extended state and apply the results with 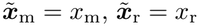, and 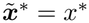. Accordingly, the costate *λ*_m_ is a scalar-valued function associated with floral tissue. For convenience, we introduce the rescaled costate 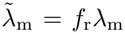. Using eqs. (40) and (41) with 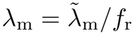, the costate satisfies

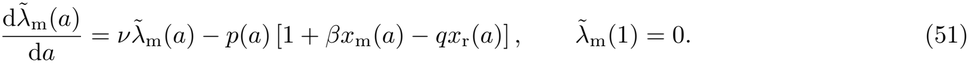

Using eq. (50) in Result 5, the selection gradient on investment at age *a* is therefore

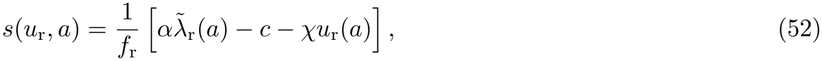

where 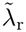 is the resident costate, obtained by setting *x*_m_ = *x*_r_ and 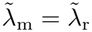 in eq. (51).

Increasing investment at age *a* produces *α* additional units of floral tissue per unit investment, whose future reproductive value is 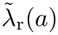, but entails the marginal investment cost *c* + *χu*_r_(*a*). As in the preceding example, selection balances a marginal benefit against a marginal cost. Here, the benefit is the future reproductive value of the floral tissue produced at age *a*, captured by the costate 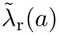.

For an interior singular growth schedule, directional selection vanishes at every age, giving

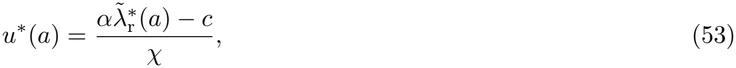

and together with the state and costate equations yields

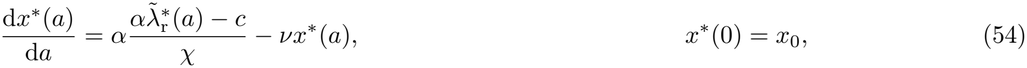

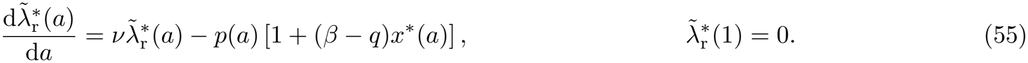

Equations (54) and (55) can be solved numerically to obtain the singular investment schedule *u*^*\**^(*a*) (using eq. (53)) and the resulting floral tissue trajectory *x*^*\**^(*a*).

Next, we assume that the solution satisfies *u*^*\**^(*a*) *>* 0 throughout the season, so that *u*^*\**^ is entirely interior. To compute the singular strategy numerically, we consider a seasonal environment in which pollinator abundance *p*(*a*) peaks at mid-season (*a* = 0.5). In the previous example, which neglects developmental constraints, floral investment directly peaks at the time of highest pollinator abundance. Here, individuals invest heavily early in the season to produce floral tissue. Floral tissue then accumulates through time and reaches its maximum after pollinator abundance has already peaked (see fig. 2). This illustrates how developmental constraints can substantially alter the timing of flowering.

**Figure 2:**
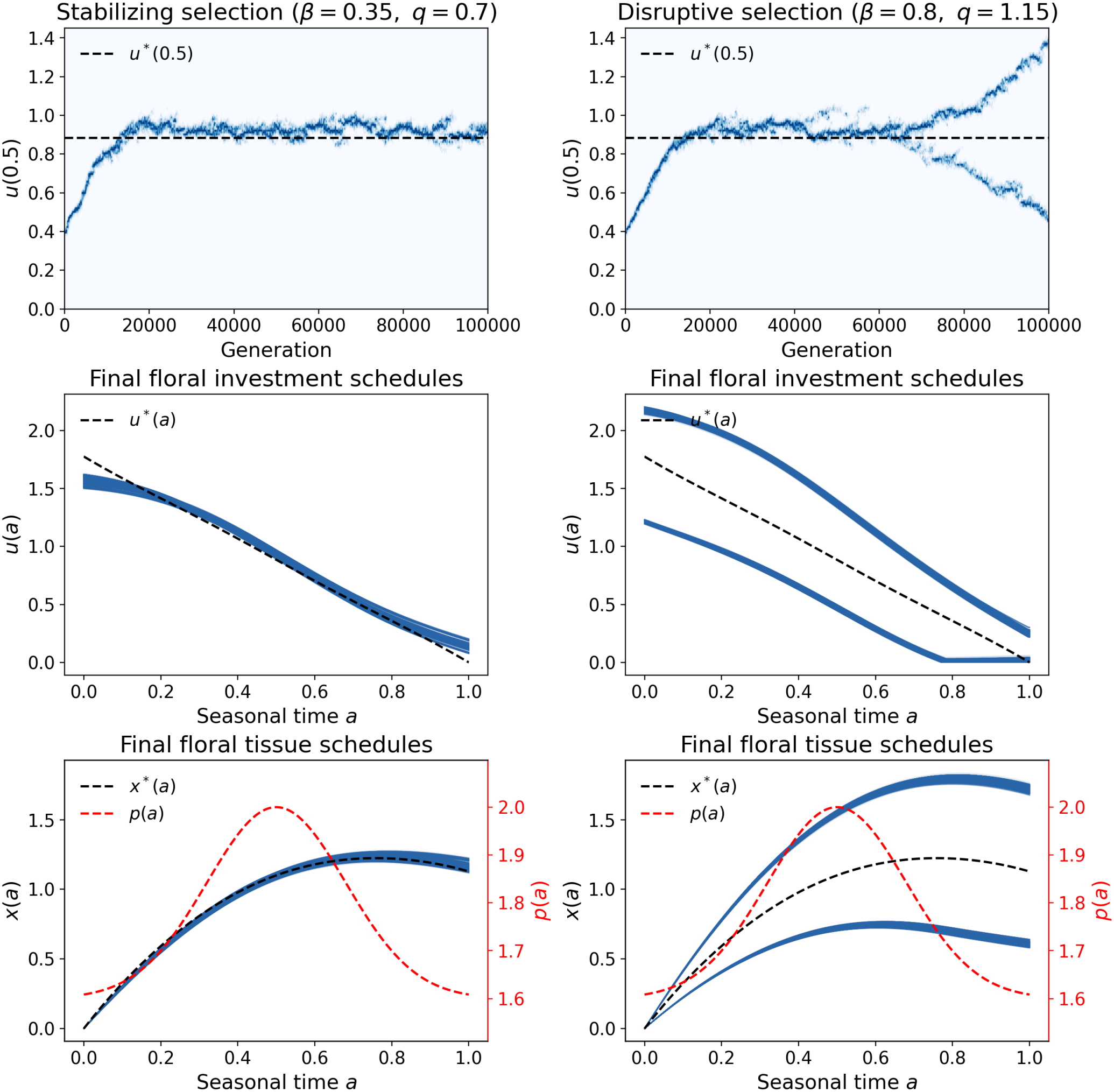
Individual-based evolution of floral growth schedules under stabilizing and disruptive selection, when floral tissue is a state variable produced by an age-dependent control. Top panels show the distribution of mid-season investment in floral growth, *u*(0.5), through time, with darker blue indicating greater density. Middle panels show the floral investment (control) schedules *u*(*a*) expressed by all individuals in the final population, with the singular control *u*^*\**^(*a*) dashed. Bottom panels show the resulting floral-tissue (state) trajectories *x*(*a*), with the singular state *x*^*\**^(*a*) dashed in black and pollinator availability *p*(*a*) shown on a separate right-hand axis in red. In the stabilizing case, we use *β* = 0.35 and *q* = 0.70, whereas in the disruptive case we use *β* = 0.80 and *q* = 1.15. Common parameters were *α* = 2.00, *v* = 0.70, *x*_0_ = 0, *c* = 0, *χ* = 1.00, *b*_0_ = 2.00, and 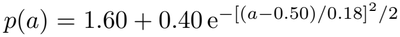. Simulation details are provided in Appendix C.2.

#### 3.3.3 Convergence stability

To determine whether selection brings nearby resident growth schedules toward *u*^*\**^, we apply Result 6 directly to the Hamiltonian in eq. (50). Since *∂*^2^*H/∂u*_m_*∂x*_m_ = 0, and using the singularity condition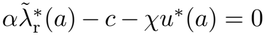, the Jacobian kernel reduces to

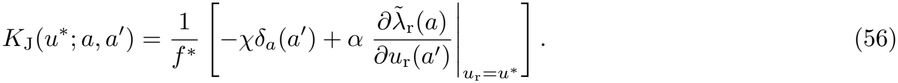

It remains to determine how the resident costate changes when resident investment changes. Differentiating the resident costate equation (51) with respect to *u*_r_(*a*^*′*^) therefore gives

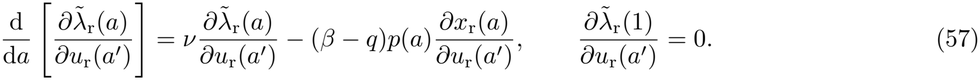

We need to express the change in floral tissue at a later age *a* due to resident investment at age *a*^*′*^, *∂x*_r_(*a*)*/∂u*_r_(*a*^*′*^). Solving eq. (47) gives

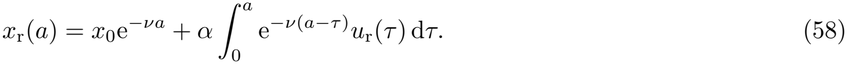

Taking the functional derivative with respect to *u*_r_(*a*^*′*^) therefore gives

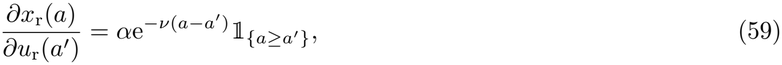

since investment at age *a* affects floral tissue only for *a ≥ a*^*′*^.

We next determine how the costate at age *a* changes with resident investment at age *a*^*′*^. Let

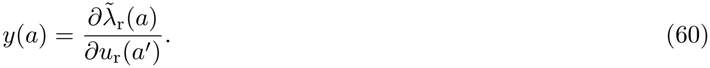

Using eq. (59), the costate sensitivity equation becomes

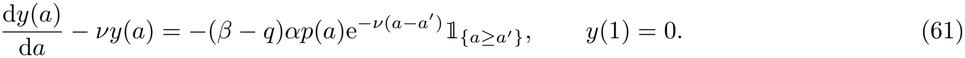

Multiplying by e^*−νa*^, and noting that

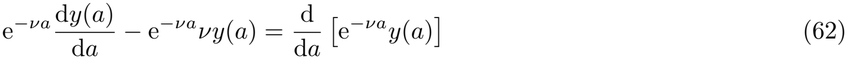

gives

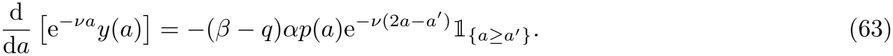

Integrating from *a* to 1 and using *y*(1) = 0 gives

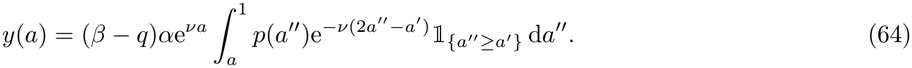

Hence,

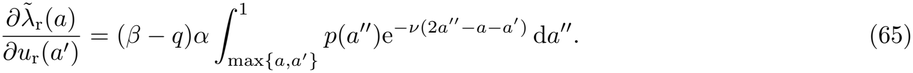

Substituting this expression into eq. (56) gives the Jacobian kernel

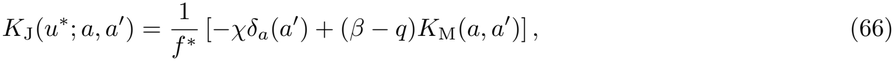

where

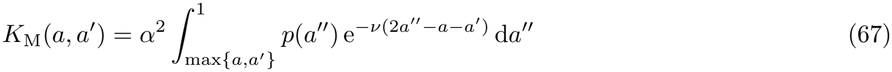

collects the state-mediated effects. The Jacobian kernel therefore separates two pathways: the cost of investment, which acts at a single age and appears through *δ*_*a*_(*a*^*′*^), and the return on floral tissue, which couples all ages through *K*_M_.

It follows that the quadratic form associated with the Jacobian is

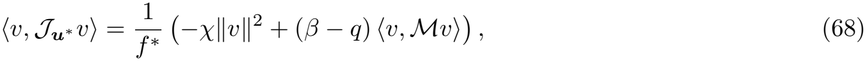

where *ℳ* denotes the integral operator with kernel *K*_M_.

The first term is always restoring because the convex cost of floral investment opposes deviations from the singular growth schedule. The second term captures how deviations alter floral tissue and thereby future reproductive returns, and can be restoring or destabilizing.

When *β ≤ q*, this state-mediated effect is restoring or vanishes because *ℳ* is positive semidefinite, so

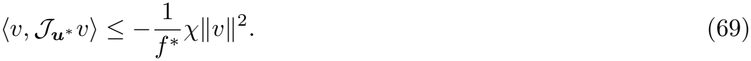

Thus, whenever *χ >* 0, 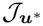 is strongly negative definite, and Result 2 implies that the singular schedule is convergence stable for any positive definite genetic variance–covariance operator.

When *β > q*, the state-mediated effect can instead oppose the direct restoring effect of the investment cost. To quantify its maximum strength, define

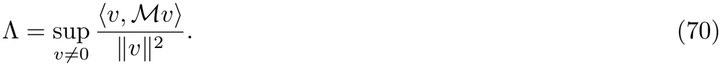

The spectral theorem for compact self-adjoint operators (Brezis, 2011, Theorem 6.11) implies that *L*^2^(*A*) admits an orthonormal basis of eigenfunctions of *ℳ*. Because *ℳ* is self-adjoint and positive, its eigenvalues are real and non-negative, and its largest eigenvalue is attained by a corresponding eigenfunction. By Theorems 4.20 and 4.24 of Arbogast and Bona (2025), Λ is the largest eigenvalue of *ℳ*. Numerically, it can be obtained by discretizing *K*_M_ on an age grid and computing the leading eigenvalue of the resulting matrix.

It follows that

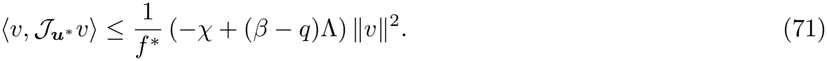

Therefore, if (*β − q*)Λ *< χ*, then 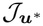 is strongly negative definite, and Result 2 implies that the singular schedule is convergence stable for any positive definite genetic variance–covariance operator.

If (*β − q*)Λ *> χ*, let *v*_Λ_ denote an eigenfunction of *ℳ* associated with Λ. Then

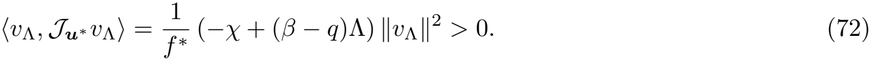

Thus, 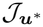 is not negative semidefinite, and Result 2 implies that there exists a positive definite genetic variance– covariance operator for which *u*^*\**^ is convergence unstable. When (*β − q*)Λ = *χ*, we cannot conclude.

#### 3.3.4 Uninvadability and polymorphism

Finally, we determine whether nearby mutant growth schedules can invade once the population has reached *u*^*\**^. To apply Result 7, we first calculate the sensitivity of the mutant costate to changes in mutant investment. Following the same procedure used to calculate the resident costate sensitivity, we obtain

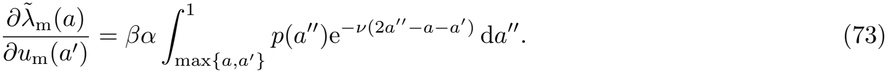

Applying Result 7 using eqs. (50) and (73) gives the Hessian kernel

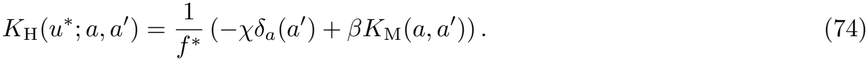

Consequently, for any functional displacement *v*,

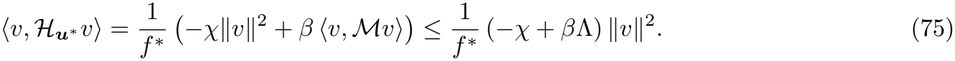

Hence, if *β*Λ *< χ*, the Hessian is strongly negative definite, and Result 3 implies that *u*^*\**^ is locally uninvadable. Otherwise, if *β*Λ *≥ χ*,

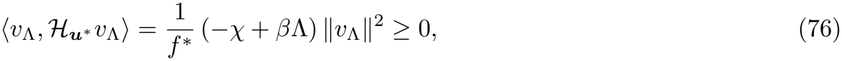

so that 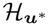 is not strongly negative definite. Result 4 then implies that there exist mutually invadable control schedules arbitrarily close to *u*^*\**^, making the singular schedule a candidate evolutionary branching point.

We numerically computed Λ and chose two parameter sets satisfying the convergence-stability condition (*β − q*)Λ *< χ*. The first also satisfies *β*Λ *< χ* and is therefore locally uninvadable, whereas the second satisfies *β*Λ *≥ χ* and is therefore a candidate branching point. We then ran individual-based simulations for these two cases to examine the corresponding evolutionary dynamics (see fig. 2). Individual-based simulations recover both evolutionary outcomes: stabilizing selection in one case and disruptive selection leading to evolutionary branching in the other. After branching, the two lineages express distinct control schedules and consequently follow different state trajectories.

## 4 Discussion

We analyzed the local gradual evolution of function-valued traits belonging to Hilbert spaces, thus covering both finite- and infinite-dimensional traits. Directional selection on a function-valued trait can be characterized by a selection gradient function, which quantifies the fitness consequences of small deviations from the resident trait function (Gomulkiewicz and Beder, 1996; Dieckmann et al., 2006; Parvinen et al., 2006, 2013). We then derived the criteria for classifying singular functions (section 2.5), at which directional selection vanishes in the interior of the trait space. This is based on the properties of two operators. The Jacobian of the selection gradient function provides conditions for the local convergence stability of singular functions, whereas the Hessian of invasion fitness provides conditions for their local uninvadability. Moreover, when the Jacobian is strongly negative definite but the Hessian is not, mutually invadable trait functions exist arbitrarily close to the singular function. These results extend the classical criteria for classifying scalar and multidimensional singular traits (e.g., Geritz et al., 1998; Leimar, 2009) to function-valued traits, and further clarify those stated for function-valued traits by Dieckmann et al. (2006, Sect. 4). Our main result thus shows that, provided the stability of the linearized evolutionary dynamics around a singular function is understood in the sense of strong stability (eq. (16)), results from finite dimensions carry over to the infinite-dimensional case. A key difference in infinite dimensions, however, is that the sufficient conditions for convergence stability and uninvadability require the Jacobian and Hessian to be strongly negative definite, not merely negative definite; namely, selection must oppose every displacement with strength bounded away from zero, because it can otherwise become arbitrarily weak along some displacements. Together, our results allow us to distinguish evolutionary outcomes in which populations remain monomorphic at a singular function from those in which evolutionary branching is possible.

Function-valued traits include life-history traits whose expression depends on age. In this case, a trait expressed at one age may affect reproduction immediately, but also alter a biological state, such as body size, physiological damage, or acquired knowledge, with consequences for reproduction later in life. These effects complicate the evolutionary analysis, and tools from optimal-control theory have long been used to disentangle them and study the evolution of life-history traits (e.g., León, 1976; Iwasa and Roughgarden, 1984; Perrin, 1992; Day and Taylor, 2000; Mangel, 2008). In particular, they allow the selection gradient to be expressed as the sum of direct effects on current reproduction and indirect effects on future reproduction through changes in survival and biological states (Metz et al., 2016; Avila et al., 2021). We extend this approach by deriving a representation for the corresponding Jacobian and Hessian operators (eqs. (45)–(46)), which account for how changes in trait expression at different ages affect state trajectories and reproductive success. These results provide criteria for determining when singular life-history traits are convergence stable, uninvadable, or allow evolutionary diversification.

Our analysis, however, does not allow us to formalize the full diversity of life-history schedules observed in nature and could be extended in at least five biologically relevant directions. First, our life-history analysis considers only open-loop controls, where trait expression follows a fixed age-dependent schedule, ***u***(*a*). Yet investment decisions may also depend on an individual’s current biological state, such as body size or energy reserves. Extending our analysis to such closed-loop controls, where states affect subsequent trait expression, ***u***(*a*, ***x***(*a*)), would require accounting for this feedback when deriving the selection gradient function, the Jacobian, and the Hessian operators (Avila et al., 2021). Second, we assume that individuals begin life with a fixed biological state, independently of their parents. Yet parental effects, including resource provisioning, hormonal and epigenetic influences, immune factor transmission, and social learning, can make offspring biological states depend on those of their parents (Badyaev and Uller, 2009; Bonduriansky and Day, 2009). Incorporating these effects would introduce intergenerational state dynamics, making the evolutionary analysis more complex (Maisonneuve and Lehmann, 2026). Third, our second-order results concern interior singular functions at which invasion fitness has curvature in the trait expressed at each age. In many classical life-history models, however, the trait enters the vital rates linearly (e.g., León, 1976; Iwasa and Roughgarden, 1984; Perrin, 1992). Candidate life histories then often lie on the bounds of the admissible set (bang-bang solutions), but may also contain interior stretches along which fitness has no curvature in the trait (singular arcs) (Bryson and Ho, 1975). Our second-order criteria are inconclusive for such traits, but this does not mean that gradual evolution cannot approach them. On the bounds, directional selection alone holds the trait against the bound. Along singular arcs, by contrast, convergence and uninvadability have received little attention in life-history evolution, and characterizing them is beyond the scope of this paper. Fourth, our framework does not account for class structure, although many populations comprise distinct classes, such as males and females, social castes, or habitat structure. Weyna et al. (2026) characterized the gradual evolution of traits that remain constant throughout the life of individuals under state-dependent ontogeny in class-structured populations. When our controls are constant across age and their model is restricted to a single class, the two approaches are consistent. Combining these approaches could therefore allow us to extend our criteria for convergence stability, uninvadability, and evolutionary branching to age-dependent controls in class-structured populations. Finally, we assumed deterministic growth trajectories. Extending the model to stochastic growth trajectories and considering the evolution of stochastic controls (Malliaris and Brock, 1982) would also be interesting.

Overall, our results extend evolutionary invasion analysis to function-valued traits, providing a general approach for predicting when gradual evolution leads to stable evolutionary endpoints or creates opportunities for diversification.

## Acknowledgments

We gratefully acknowledge Charles Mullon and Adrien Blanchet for useful discussions.

## Code

All code used in this study is accessible at https://github.com/Ludovic-Maisonneuve/function-valued-traits-evolution.

## Appendix A: Results for function-valued traits

Here, we derive the results presented in the main text within the general framework of function-valued traits.

### A.1 Result 1

We prove Result 1 by showing that any function that fails the candidate conditions admits an admissible nearby mutant with invasion fitness exceeding one. Suppose that ***u***_r_ does not satisfy eq. (8) with ***u***^*\**^ = ***u***_r_. Then there exists ***v*** such that ***u***_r_ + *ϵ****v*** *∈ U*_ad_ and

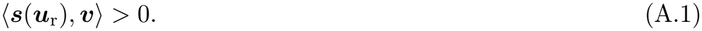

Equation (4) then gives

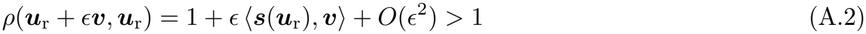

for sufficiently small *ϵ*. Thus, an admissible mutant close to ***u***_r_ can invade, and ***u***_r_ cannot be a candidate endpoint. This proves Result 1.

### A.2 Result 2

We prove Result 2 by studying how the distance between the resident trait and the singular trait changes under the evolutionary dynamics.

We assume that *G* is trace-class, positive definite, and self-adjoint. Biologically, this means that genetic variation has finite total variance and is present in every non-zero phenotypic direction. Formally, recall that a linear operator *G* with domain and range *X* is: (i) bounded if there exists a real number *c* such that *∥G****v****∥ ≤ c∥****v****∥* for all ***v*** *∈ X* (Kreyszig, 1991, *p. 91); (ii) positive (negative) semidefinite if ⟨G****v, v****⟩ ≥* 0 (*⟨G****v, v****⟨ ≤* 0) for all ***v*** *∈ X* ; (iii) positive (negative) definite if *⟨G****v, v****⟩ ≥* 0 (*⟨G****v, v****⟩ ≤* 0) for all ***v*** *∈ X* and *⟨G****v, v****⟩ >* 0 (*⟨G****v, v****⟩ <* 0) for ***v*** ≠ **0**, and (iv) strongly positive (negative) definite if for all ***v*** *∈ X, ⟨G****v, v****⟩ ≥ c∥****v****∥*^2^ (*⟨G****v, v****⟩ ≤ −c∥****v****∥*^2^) for some *c >* 0 (Phat and Kiet, 2002, p. 156).

The adjoint *G*^*†*^ of the operator *G* is the operator that satisfies *⟨G****v, µ****⟩* = *⟨****v***, *G*^*†*^***µ****⟩* for all ***v*** *∈ X* and ***µ*** *∈ X* (Kreyszig, 1991, *p. 196, note that if G* is a matrix and has real entries then the adjoint and the transpose are the same) and *G* is self-adjoint if *G* = *G*^*†*^ (Kreyszig, 1991, *p. 201). Let (T* (*t*))_*t≥*0_ be a family of bounded linear operators on *X* . It is called a strongly continuous (or *C*_0_) semigroup if

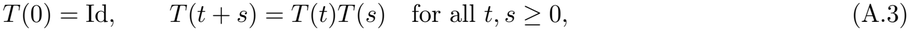

where Id denotes the identity operator on *X* and, for every ***v*** *∈ X*, the map *t ↦ T* (*t*)***v*** is continuous (Schnaubelt, 2011, Definition 1.1, p. 1). When applied to a linear operator *T, ∥ · ∥* denotes the operator norm, defined by

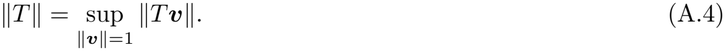

The semigroup is bounded if

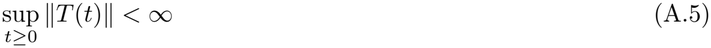

(Schnaubelt, 2011, Definition 1.5, p. 3). The semigroup is holomorphic (or analytic) if it admits an extension *T* (*z*) to a complex sector around the positive real axis such that *z ↦ T* (*z*) is complex differentiable (Schnaubelt, 2011, Definition 2.24, p. 63). Finally, the semigroup is strongly stable if

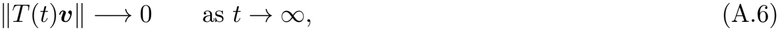

for every ***v*** *∈ X* (Eisner, 2010, *Definition 3.1, p. 100, Engel and Nagel, 2000, p. 296)*.

#### A.2.1 Local stability of u^*^

Define the displacement from the singular function as

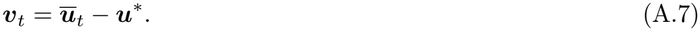

Because ***u***^*\**^ is fixed, substituting eq. (A.7) into eq. (15) gives

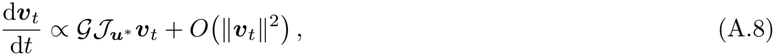

where 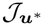 is the Jacobian operator of the selection gradient at ***u***^*\**^, defined through its kernel by

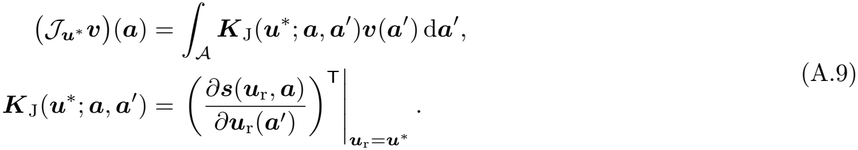

After rescaling evolutionary time to absorb the positive proportionality constant, the linearized dynamics around ***u***^*\**^ read

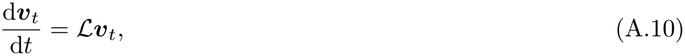

where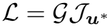.

Assume, as in Result 2, that 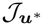 is strongly negative definite, that is, there exists a constant *c*_J_ *>* 0 such that

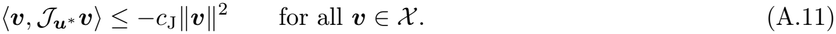

We say that the trait value ***u***^*\**^ is convergence stable if the linearized dynamics around ***u***^*\**^ given in eq. (A.10) are strongly stable, that is, if *∥****v***_*t*_*∥ −→* 0 as *t → ∞*. Let (*T* (*t*))_*t≥*0_ denote the *C*_0_-semigroup generated by *ℒ*, so that

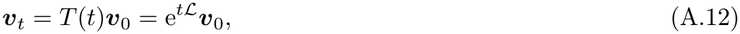

where ***v***_0_ is the displacement from ***u***^*\**^ at the reference time from which the local dynamics around ***u***^*\**^ are considered (see Engel and Nagel, 2000, Chapter I, Theorem 3.7; Chapter II, Proposition 6.4). To establish strong stability, we apply Corollary 3.17 of Eisner (2010), which states that *T* is strongly stable if the following two conditions are satisfied: (i) 0 is not an eigenvalue of *ℒ*^*†*^ and (ii) *T* is a bounded holomorphic *C*_0_-semigroup.

##### First condition

We first show that 0 is not an eigenvalue of *ℒ*^*†*^, or equivalently that

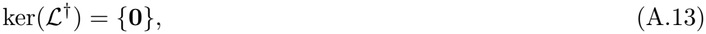

where ker denotes the kernel of an operator, i.e., the set of vectors that it maps to zero. Since 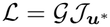 and *G* is self-adjoint,

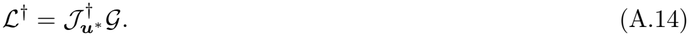

Both operators appearing in this product have trivial kernels. First, the positive definiteness of *G* implies

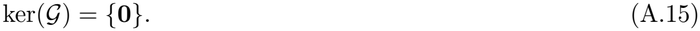

Second, by the definition of the adjoint and strong negative definiteness of 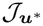 (Kreyszig, 1991, *p. 196)*,

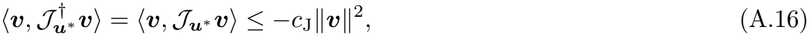

so 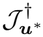 is strongly negative definite and therefore

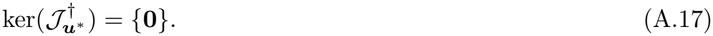

Now let ***v*** *∈* ker(*ℒ*^*†*^). Then

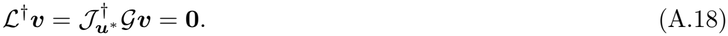

Since ker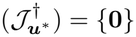, this implies *G****v*** = **0**. Since ker(*G*) = *{***0***}*, it follows that ***v*** = **0**. Hence

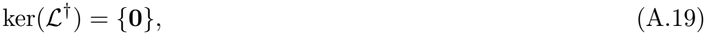

and therefore 0 is not an eigenvalue of *ℒ*^*†*^.

##### Second condition

It remains to show that (*T* (*t*))_*t≥*0_ is a bounded holomorphic *C*_0_-semigroup. By Theorem 2.25 of Schnaubelt (2011), condition (b) of that theorem is equivalent to condition (e), which states that *T* is a bounded holomorphic *C*_0_-semigroup. It is therefore sufficient to establish condition (b), that is, for every *λ ∈* ℂ satisfying ℛ (*λ*) *>* 0, where ℛ (*z*) denotes the real part of a complex number *z*, the operator *λ*Id *−ℒ* is bijective and its inverse satisfies

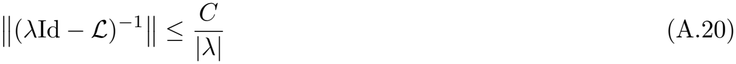

for some constant *C >* 0 independent of *λ*. The theorem also requires *ℒ* to be closed and densely defined. These conditions are automatically satisfied here because *ℒ* is a bounded operator defined on all of *X* (Schnaubelt, 2011, *Example 1.15(a)): G* is bounded because it is trace-class, and 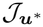 is bounded because it is the Fréchet derivative at ***u***^*\**^ of the selection gradient ***s***(***u***_r_), which is Fréchet differentiable since *ρ* is three times continuously Fréchet differentiable.

Because Theorem 2.25 of Schnaubelt (2011) requires the condition for complex values of *λ*, we consider the complexification *X*_ℂ_ = *{****x*** + *i****y*** : ***x, y*** *∈ X}* of *X*, equipped with the inner product

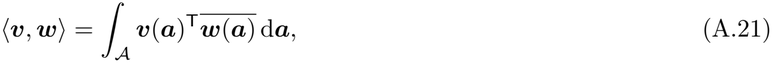

which is linear in the first argument and conjugate-linear in the second. Indeed, when *λ* is complex, *λ*Id *− ℒ* does not in general map the real space *X* into itself. The evolutionary dynamics remain real; we use complexification only for this spectral argument.

We now verify the conditions required by Theorem 2.25 of Schnaubelt (2011). Our goal is therefore to show that, for every *λ* with (*λ*) *>* 0, *λ*Id *−ℒ* is bijective and that its inverse is bounded by *C/*|*λ*|. To do so, we factorize *λ*Id *− ℒ* into two bijective operators. We first show that 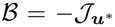 is invertible, and therefore bijective, and use this to write

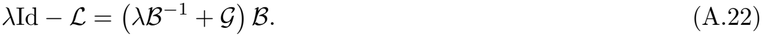

By Corollary 5.49 of Schnaubelt (2025, p. 114), *ℬ* is invertible and thus bijective if *∥ ℬ****v****∥ ≥ c*_J_*∥****v****∥* for all ***v*** *∈ X*_ℂ_ and *ℬ*^*†*^ is injective. We first show the first condition. Let ***v*** = ***x*** + *i****y*** *∈ X*_ℂ_, where ***x, y*** *∈ X* . By Cauchy–Schwarz,

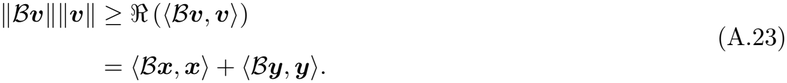

Since 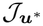 is strongly negative definite, there exists *c*_J_ such that

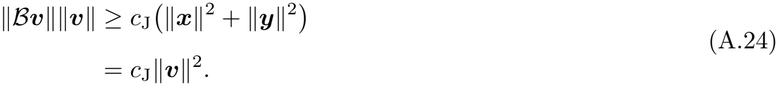

Thus

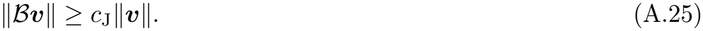

We now show that *ℬ*^*†*^ is injective. By the definition of the adjoint, for all ***v*** *∈ X*_ℂ_,

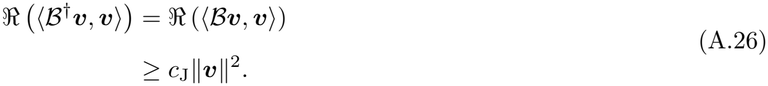

Thus, *ℬ*^*†*^***v*** = **0** implies ***v*** = **0**, so *ℬ*^*†*^ is injective. Thus, both conditions of Corollary 5.49 of Schnaubelt (2025, p. 114) are satisfied, so *ℬ* is invertible and hence bijective.

It remains to show that

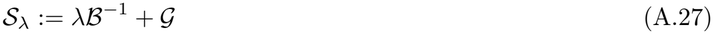

is bijective. By Corollary 5.49 of Schnaubelt (2025, p. 114), it is sufficient to prove that

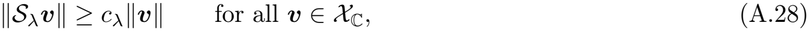

for some *c*_*λ*_ *>* 0, and that 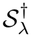 is injective.

Both properties will follow if we can find an operator *C*_*λ*_ such that

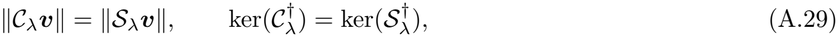

and

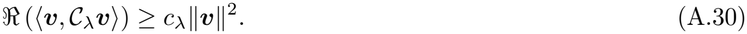

Indeed, by Cauchy–Schwarz,

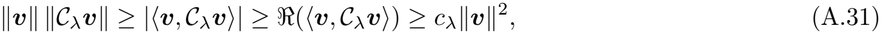

so that

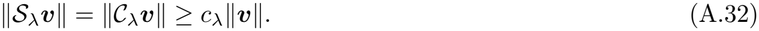

Moreover, by the definition of the adjoint,

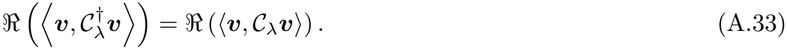

Therefore, eq. (A.30) also gives

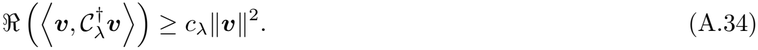

If 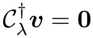, the left-hand side is zero, and hence

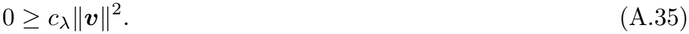

Since *c*_*λ*_ *>* 0, this implies ***v*** = **0**. Thus, 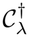 is injective.

We now construct such an operator. Since ℛ (*λ*) *>* 0, write

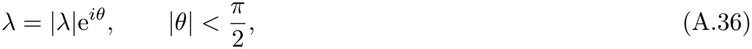

and define

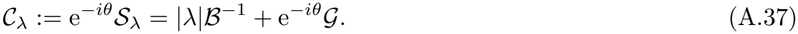

Since |e^*−iθ*^| = 1,

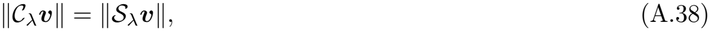

and since

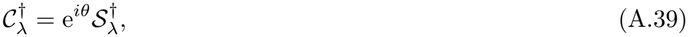

the two adjoints have the same kernel.

Thus, it only remains to prove eq. (A.30). Since *G* is self-adjoint, we have

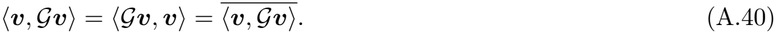

Hence, (***v***, *G****v***) is real, even for complex ***v***. Using this property and the conjugate-linearity of the inner product in its second argument, eq. (A.37) gives

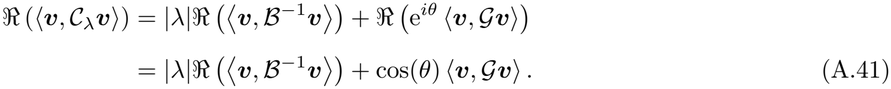

Since ℛ (*λ*) *>* 0, we have cos(*θ*) *>* 0. Moreover, writing ***v*** = ***x*** + *i****y***, where ***x, y*** *∈ X*, the self-adjointness and positive definiteness of *G* imply

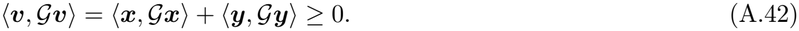

Therefore, the second term in eq. (A.41) is non-negative. To obtain the required strictly positive lower bound for *C*_*λ*_, it is therefore sufficient to show that, for some *α >* 0,

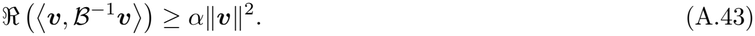

We now establish this bound. Let ***v*** = ***x*** + *i****y*** *∈ X*_ℂ_, where ***x, y*** *∈ X* . Since 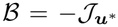 and 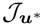 is strongly negative definite, we have

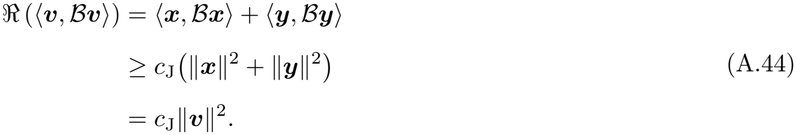

For any ***w*** *∈ X*_ℂ_, let ***v*** = *ℬ*^*−*1^***w***, so that ***w*** = *ℬ****v***. Then

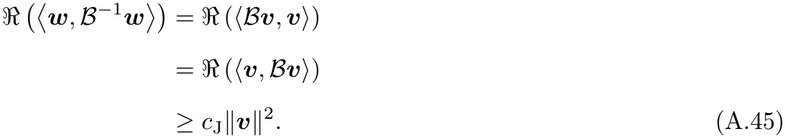

Moreover,

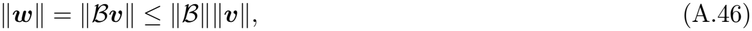

and hence

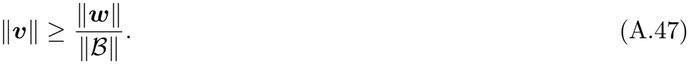

Therefore,

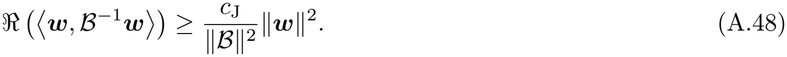

Thus eq. (A.43) holds with

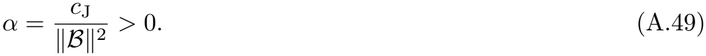

Substituting this bound into eq. (A.41) yields

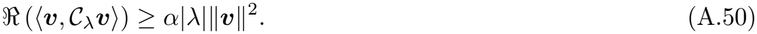

Hence eq. (A.30) holds with *c*_*λ*_ = *α*|*λ*| *>* 0.

Altogether, this shows that *S*_*λ*_ is invertible and therefore bijective.

It remains to establish the required bound on its inverse. Let ***w*** *∈ X*_ℂ_. Since *S*_*λ*_ is bijective, there exists a unique ***v*** *∈ X*_ℂ_ such that ***w*** = *S*_*λ*_***v***. Equivalently,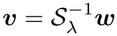. We have

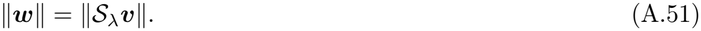

Moreover, since *c*_*λ*_ = *α*|*λ*|, eq. (A.28) gives

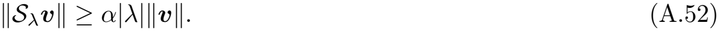

This implies that

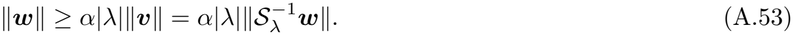

Therefore,

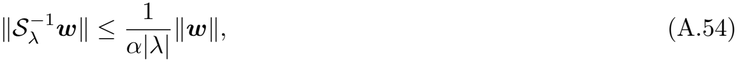

and hence

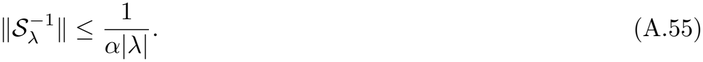

Since

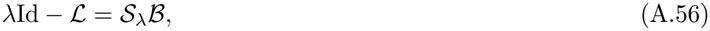

and both factors are bijective, *λ*Id *− ℒ* is bijective and

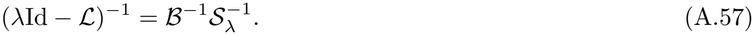

Using

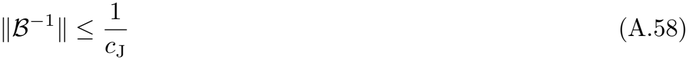

and

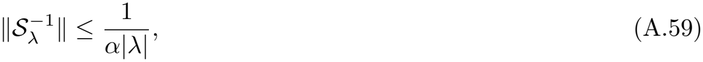

we obtain

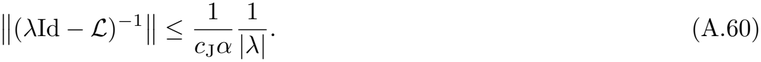

Thus condition (b) of Theorem 2.25 of Schnaubelt (2011) is satisfied. Therefore, the semigroup *T* generated by *ℒ* is a bounded holomorphic *C*_0_-semigroup.

We have also shown above that

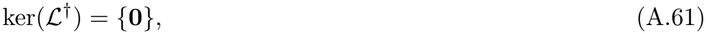

so that 0 is not an eigenvalue of *ℒ*^*†*^. Corollary 3.17 of Eisner (2010) therefore implies that (*T* (*t*))_*t≥*0_ is strongly stable. Consequently, for every initial displacement ***v***_0_ *∈ X*,

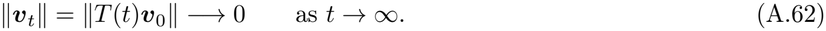

Hence the singular function ***u***^*\**^ is convergence stable.

Finally, the antisymmetric part of 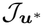 does not contribute to its quadratic form:

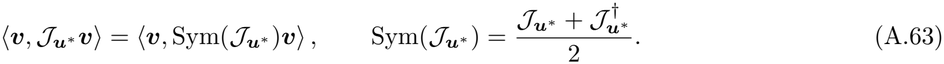

Because Sym 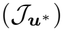 is self-adjoint, Theorems 9.2-1 and 9.2-3 of Kreyszig (1991, pp. 465–467) imply that

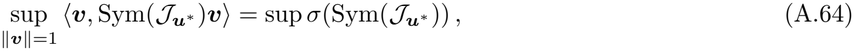

where *σ* denotes the spectrum of an operator (e.g., Kreyszig, 1991, p. 371). Consequently,

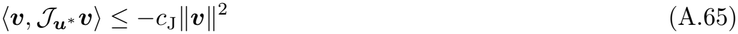

for every displacement if and only if

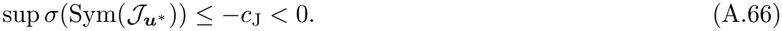

In other words, 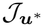 is strongly negative definite if and only if the leading spectral value of the symmetric part of 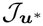 is negative, and this condition is sufficient for ***u***^*\**^ to be a local attractor of the evolutionary dynamics.

#### A.2.2 Local instability of *u*^*\**^

Our goal is to construct a genetic variance–covariance operator under which ***u***^*\**^ is linearly unstable. For a genetic variance–covariance operator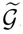, the rescaled linearized evolutionary dynamics are

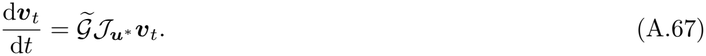

In order to show the instability of these dynamics, it is sufficient to find a non-zero direction ***v*** and an admissible operator 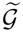 such that

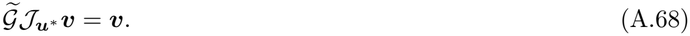

Indeed, ***v*** is then an eigenfunction of the linearized evolutionary operator with eigenvalue 1 *>* 0, so a perturbation initially along ***v*** grows as ***v***_*t*_ = e^*t*^***v***.

We now seek a direction ***v*** for which such an operator can be constructed. Because 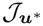 is not negative semidefinite, there exists a non-zero direction ***v***_0_ *∈ X* such that

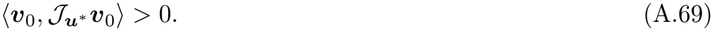

The direction ***v***_0_ does not necessarily define an admissible trait perturbation, because when the argument ***a*** varies continuously, it need not be piecewise continuous. Since 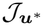 is bounded, the quadratic form 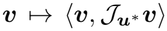 is continuous, and piecewise-continuous functions are dense in *X* (Rudin, 1974, Theorem 3.14). Hence, ***v***_0_ can be approximated by a piecewise-continuous direction ***v*** satisfying the same strict inequality. When *A* is discrete, we simply take ***v*** = ***v***_0_. In both cases, because ***u***^*\**^ is interior, ***u***^*\**^ + *ϵ****v*** *∈ U*_ad_ for sufficiently small *ϵ >* 0.

To construct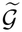, let

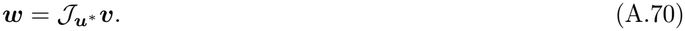

The target in eq. (A.68) is therefore equivalent to

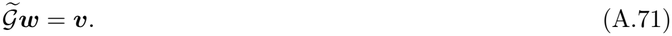

Since *X* admits a finite or countable orthonormal basis (Brezis, 2011, Theorem 4.13), (Reed and Simon, 1980, Theorem II.7), so does its closed subspace orthogonal to ***w***. Let (*e*_*n*_)_*n∈I*_ denote such a basis, with *I* = *{*1, …,*N; }* or *I* = *ℕ*. Define

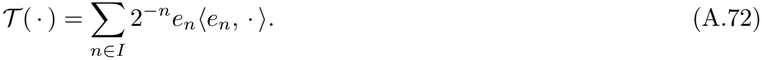

We define

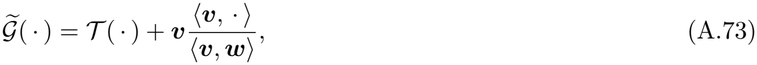

where

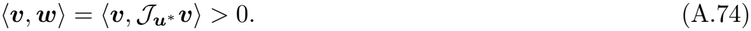

Applying this operator to ***w*** gives

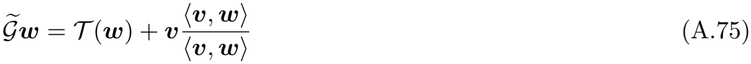

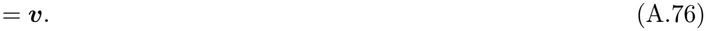

Since 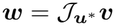, we therefore have

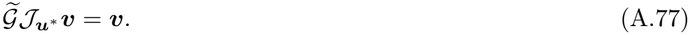

Hence ***v*** is an eigenfunction of the linearized evolutionary operator with positive eigenvalue 1.

It remains to establish that 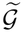 is positive definite, self-adjoint and trace-class.

For any ***h*** *∈ X*, eq. (A.73) gives

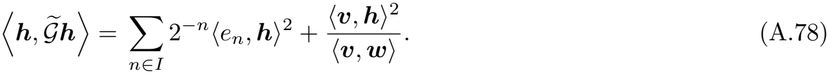

The first term is non-negative and vanishes only when ***h*** is proportional to ***w***. The second term is also non-negative because (***v, w****) >* 0. Hence

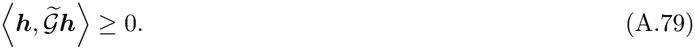

Moreover, equality can hold only when ***h*** = **0**. Indeed, equality requires the first term to vanish, which implies that ***h*** is proportional to ***w***, say

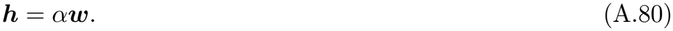

It also requires the second term to vanish, so that

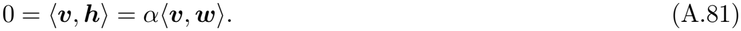

Since (***v, w****) >* 0, this implies *α* = 0, and therefore ***h*** = **0**. Thus

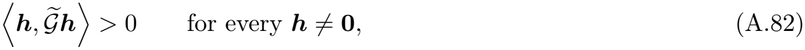

so 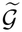 is positive definite.

It remains to show that 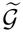 is self-adjoint. Let ***h, g*** *∈ X* . Applying eq. (A.73) and expanding gives

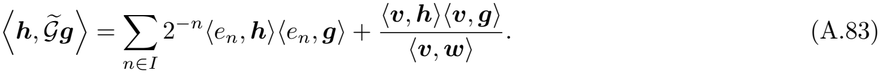

Both terms are unchanged when ***h*** and ***g*** are exchanged, so

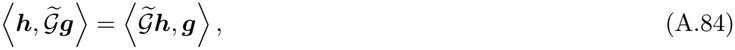

which establishes self-adjointness.

Finally, *T* has eigenvalues 2^*−n*^ on the orthogonal complement of ***w*** and 0 in the direction of ***w***, so its trace is *Σ*_*n∈I*_ 2^*−n*^. The second term of eq. (A.73) always maps functions to a multiple of ***v***. It therefore has rank one, with a single non-zero eigenvalue *∥****v****∥*^2^*/⟨****v, w****⟩*, obtained by applying it to ***v***. Its trace thus equals this eigenvalue. Thus

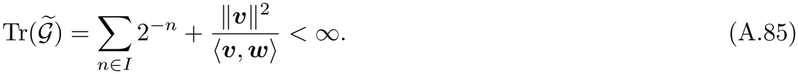

Since 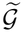 is positive and self-adjoint, we have

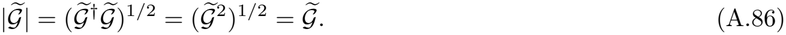

Hence, 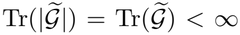 so 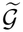 is trace-class by definition (Reed and Simon, 1980, definition following Theorem VI.18, p. 207). The operator 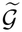 therefore satisfies all the assumptions.

Therefore, negative semidefiniteness of 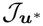 is necessary for convergence stability to hold uniformly over all self-adjoint, positive definite, trace-class genetic variance–covariance operators. If 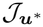 is negative semidefinite but not strongly negative definite, however, neither the sufficient stability condition nor the preceding instability construction applies.

Together, these arguments prove Result 2.

### A.3 Result 3

Here, we establish Result 3 by studying the invasion of a nearby mutant around the singular resident.

#### A.3.1 u^*^ as a local fitness maximum

The implication from the strong negative definiteness of 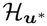 to local uninvadability stated in Result 3 is a standard second-order sufficient optimality result (Bonnans and Shapiro, 2000, Remark 3.62 and Theorem 3.63(i), pp. 188– 189). For completeness, we provide an argument directly in our notation.

Consider an admissible mutant of the form

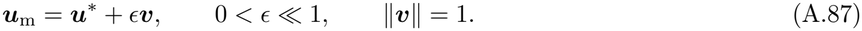

A second-order Taylor expansion with respect to the mutant trait gives

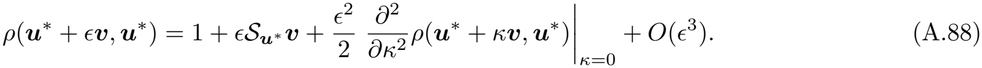

Because ***u***^*\**^ is singular, 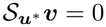 for every direction ***v***.

Using eq. (A.16) in Engel and Dreizler (2013, p. 407), the second derivative can be represented by the Hessian operator,

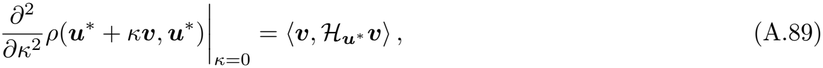

where 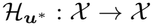 is defined by

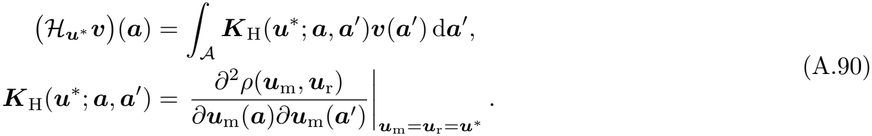

Equation (A.88) becomes

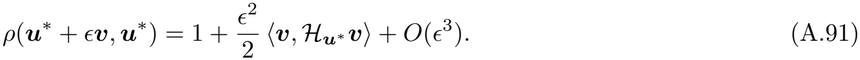

Suppose that 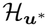 is strongly negative definite. Then there exists *c*_H_ *>* 0 such that

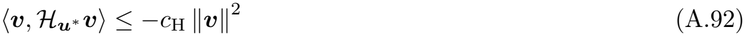

for every admissible direction ***v***.

To conclude that this negative second-order effect prevents invasion in every direction, we must show that it dominates the third-order remainder uniformly for all sufficiently small mutations. Applying Taylor’s theorem (Pathak, 2018, Theorem 3.12, pp. 146–147), the remainder in eq. (A.91) is

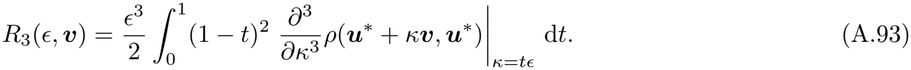

Because invasion fitness is three times continuously Fréchet differentiable, there exist *M >* 0 and *ϵ*_0_ *>* 0 such that

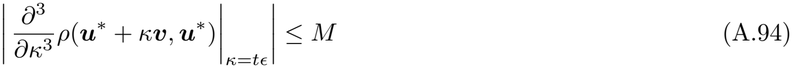

for every admissible direction ***v***, every *t ∈* [0, 1], and every 0 *< ϵ < ϵ*_0_. Therefore,

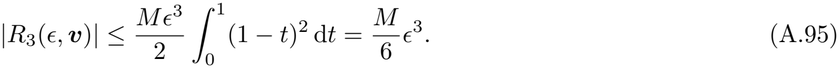

Thus, setting *C* = *M/*6, the remainder is bounded by *Cϵ*^3^ uniformly over all admissible directions. Hence

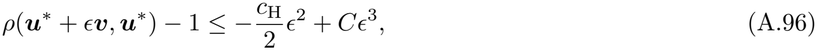

for every admissible ***v***. For all sufficiently small *ϵ >* 0, *−*(*c*_H_*/*2)*ϵ*^2^ + *Cϵ*^3^ *<* 0, and thus *ρ*(***u***^*\**^ + *ϵ****v, u***^*\**^) *−* 1 *<* 0. Therefore, no sufficiently close mutant can invade ***u***^*\**^, which proves local uninvadability.

Finally, because 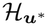 is a bounded self-adjoint operator, Theorems 9.2-1 and 9.2-3 of Kreyszig (1991, pp. 465– 467) imply that

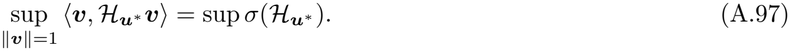

Thus, strong negative definiteness is equivalent to sup 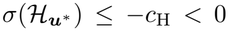, as stated in eq. (20), which proves Result 3. Then, ***u***^*\**^ is locally uninvadable if the leading spectral value of 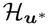 is negative.

#### A.3.2 u^*^ as locally invadable

Conversely, if 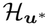 not negative semidefinite, there exists a direction ***v***_0_ *∈ X* such that

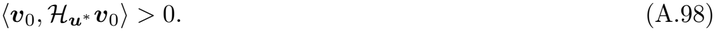

When the argument ***a*** *∈ A* varies continuously, continuity of the quadratic form and density of piecewise-continuous functions in *X* (Rudin, 1974, Theorem 3.14) allow ***v***_0_ to be approximated by a piecewise-continuous direction ***v*** satisfying the same strict inequality. When *A* is discrete, we simply take ***v*** = ***v***_0_. Because ***u***^*\**^ is interior, ***u***^*\**^+*ϵ****v*** *∈ U*_ad_ for sufficiently small *ϵ >* 0.

Since ***u***^*\**^ is singular,

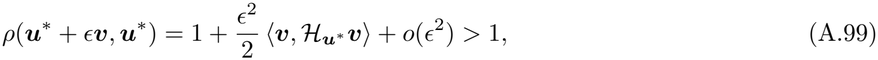

for sufficiently small *ϵ >* 0. Hence ***u***^*\**^ is locally invadable.

If 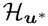 is negative semidefinite but not strongly negative definite, the second-order analysis alone is inconclusive because the quadratic term can vanish or become arbitrarily weak along some directions, so higher-order terms may determine invadability.

### A.4 Result 4

Here, we establish Result 4 by constructing two mutually invadable traits on opposite sides of ***u***^*\**^, with one trait closer to ***u***^*\**^ than the other. Let *c*_J_ *>* 0 be the constant in the convergence-stability condition (17). We assume that 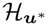 does not satisfy the sufficient condition for uninvadability given in Result 3, that is, that 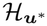 is not strongly negative definite. Therefore, for every *α ∈* (0, 1), there exists a unit direction ***v***_0_ *∈ X* such that

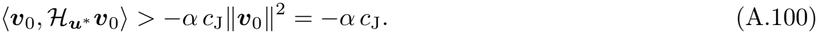

The direction ***v***_0_ does not necessarily define an admissible trait perturbation. We therefore construct a unit direction ***v*** that satisfies the same inequality and is piecewise continuous when the argument ***a*** *∈ A* varies continuously. Since the quadratic form 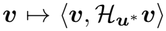 is continuous and piecewise-continuous functions are dense in *X* (Rudin, 1974, *Theorem 3.14), such a direction can be chosen arbitrarily close to* ***v***_0_. When *A* is discrete, we simply take ***v*** = ***v***_0_. Hence, in both cases,

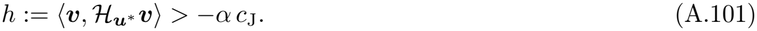

Because ***u***^*\**^ is interior, sufficiently small perturbations in both ***v*** and *−****v*** are admissible.

We keep *α* free for now and fix it below. In the same direction, convergence stability gives

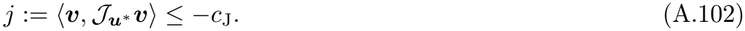

Because ***u***^*\**^ is interior, for every sufficiently small *ϵ >* 0, the two traits

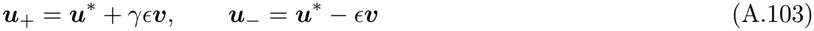

are admissible with *γ ∈* (0, 1). Because *γ <* 1, ***u***_+_ is closer to ***u***^*\**^ than ***u***_*−*_.

We first expand the invasion fitness of ***u***_+_ in a resident ***u***_*−*_ population in *ϵ* around (***u***^*\**^, ***u***^*\**^):

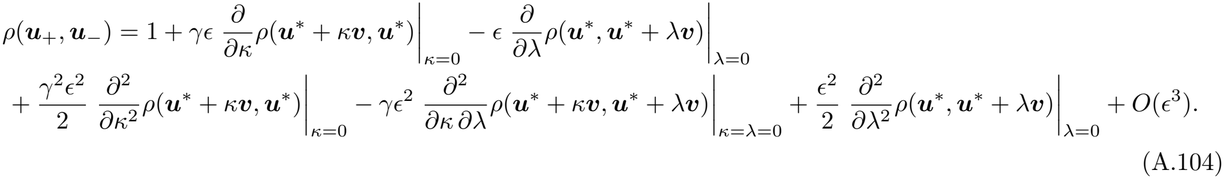

Using eq. (A.16) in Engel and Dreizler (2013, p. 407), we can write

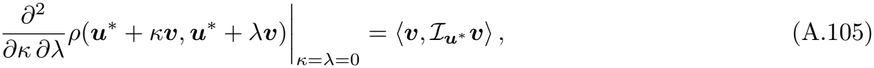

where the interaction operator 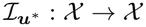 is defined by

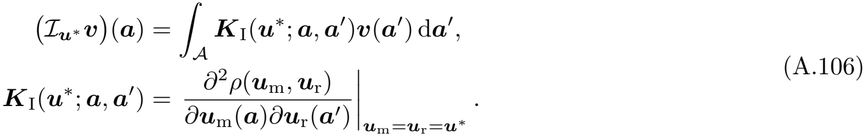

Let us now set

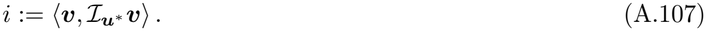

Using the notation *h, j, i*, and 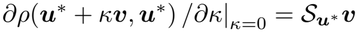, eq. (A.104) reduces to

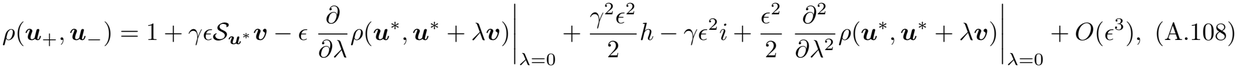

where

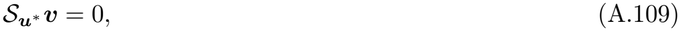

since directional selection vanishes at ***u***^*\**^.

To further simplify eq. (A.108), we use the neutrality identity

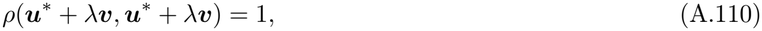

which holds for any *λ*. Differentiating once at *λ* = 0 gives

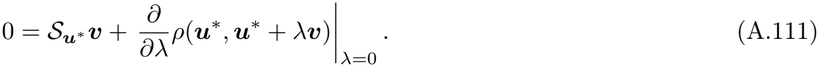

Because 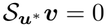, the first derivative with respect to the resident trait also vanishes:

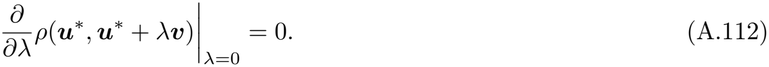

Differentiating eq. (A.110) twice at *λ* = 0 gives

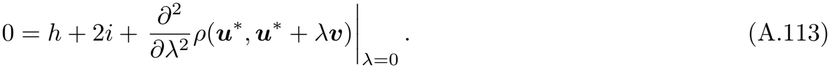

Hence, the second derivative with respect to the resident trait can be replaced by

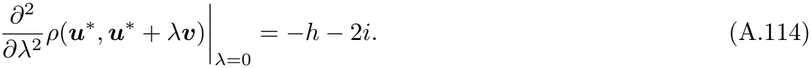

Substituting the first- and second-order diagonal identities into eq. (A.108) gives

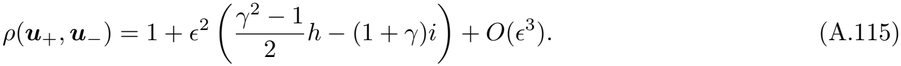

We next rearrange eq. (A.115) by relating the interaction operator to the Jacobian. Recall that the selection gradient is

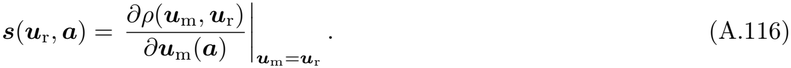

When the resident trait changes, both the mutant argument at which this derivative is evaluated and the resident argument of invasion fitness change. The chain rule therefore gives

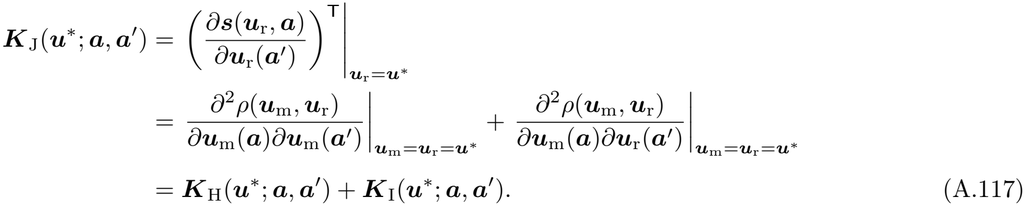

Because the kernel of 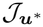 is the sum of the kernels of 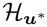 and 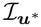, the corresponding operators satisfy

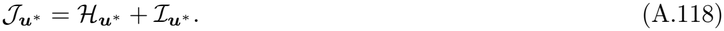

It follows that

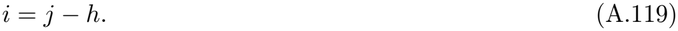

Using eq. (A.119) in eq. (A.115) yields

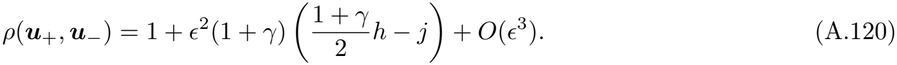

Finally, we exchange the mutant and resident traits by applying the same line of argument to invasion fitness *ρ*(***u***_*−*_, ***u***_+_). Then, a Taylor expansion gives

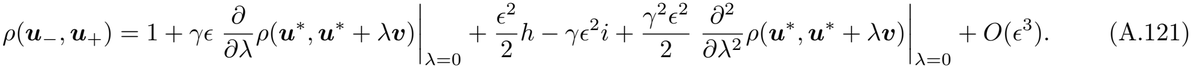

Substituting eqs. (A.112) and (A.114) into eq. (A.121) gives

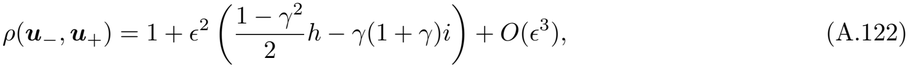

and replacing *i* by *j − h* yields

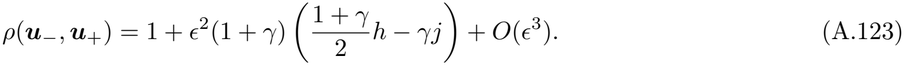

By eq. (A.101) and eq. (A.102), the bounds on *h* and *j* imply

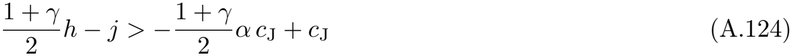

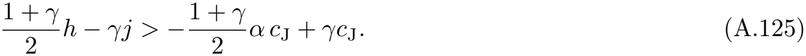

Choosing *α <* 2*γ/*(1 + *γ*) thus makes the terms in parentheses in eq. (A.120) and eq. (A.123) strictly positive, and therefore

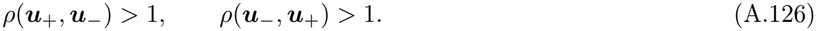

Hence ***u***_+_ and ***u***_*−*_ are mutually invadable if ***u***^*\**^ is convergence stable and 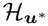 is not strongly negative definite. This proves Result 4.

## Appendix B: Results for life-history traits with state dynamics

Here, we derive the results presented in the main text within the framework of life-history traits with state dynamics, applying standard computations used in optimal control theory under the calculus of variations approach (e.g., Bryson and Ho, 1975, section 2.3, Kamien and Schwartz, 2012, section 5, Athans and Falb, 2007, section 5.7).

### B.1 Result 5

Suppose first that lifespan is finite, so that *A* = [0, *a*_f_]. To introduce the costate without changing the mutant basic reproductive number, we add to its integrand the term

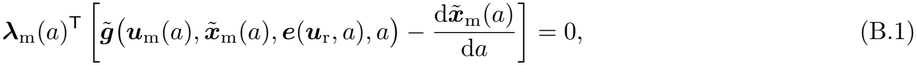

which vanishes because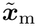 satisfies the mutant state dynamics in eq. (36). Using the definition of the Hamiltonian then gives

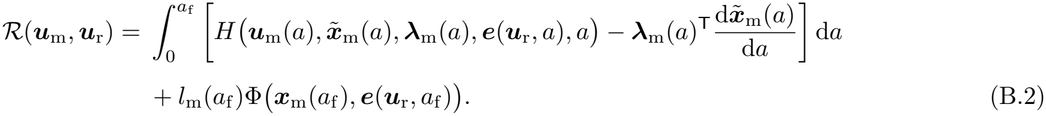

We now calculate the pointwise selection gradient. By definition,

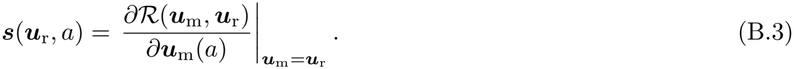

We first differentiate the mutant reproductive number in eq. (B.2) with respect to the mutant control. This gives

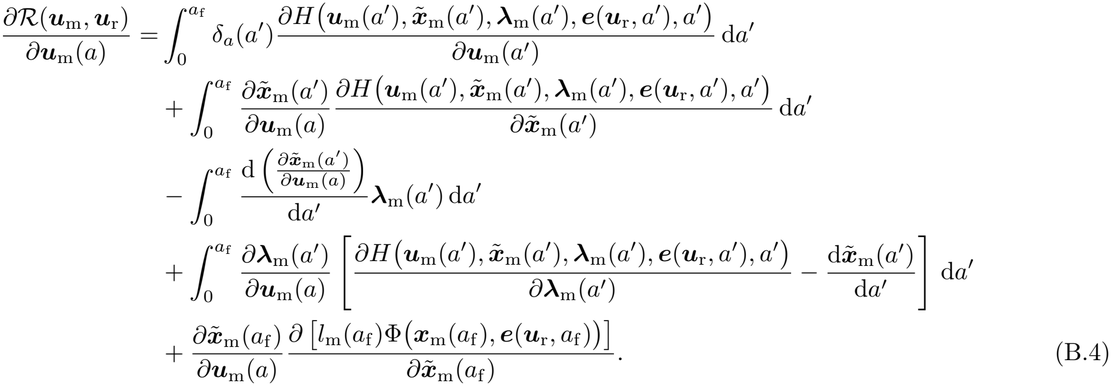

The fourth term vanishes identically because the mutant state dynamics imply

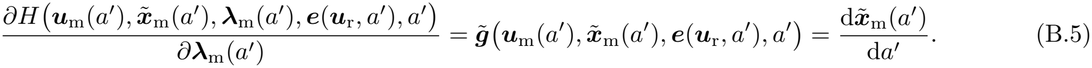

Integrating the third term by parts gives

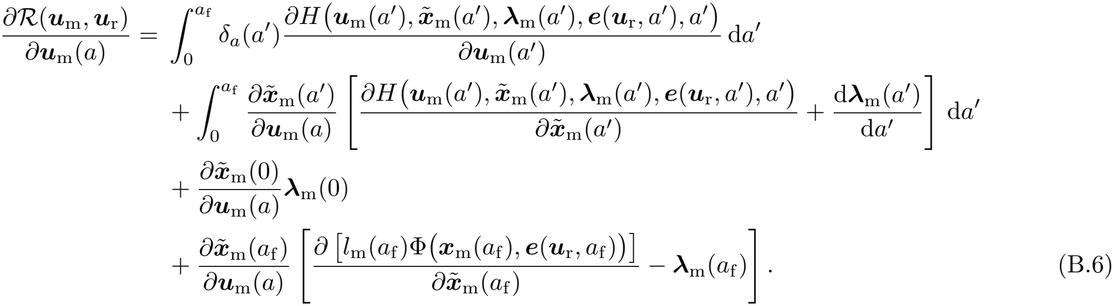

Because the initial extended state is fixed,

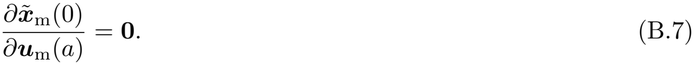

Moreover, the costate dynamics defined in eq. (40) imply

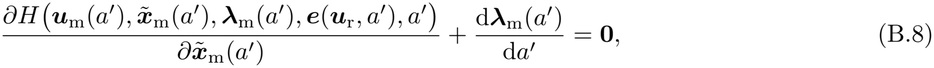

and its terminal condition, given in eq. (41), implies

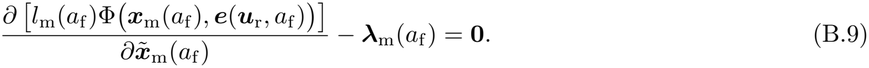

All terms involving the state sensitivity therefore vanish, leaving

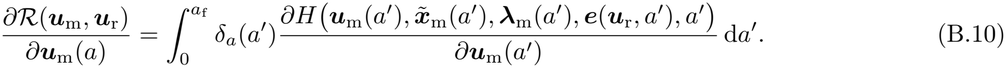

Using the defining property of the Dirac delta, we get

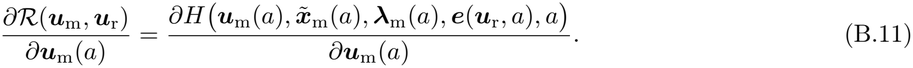

Evaluating this identity at ***u***_m_ = ***u***_r_, where 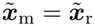 and ***λ***_m_ = ***λ***_r_, gives

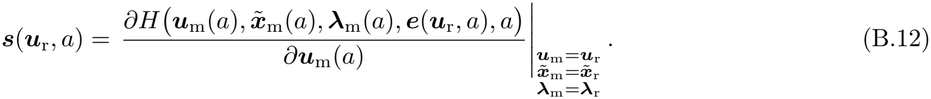

When lifespan is unbounded, we apply the same calculation over [0, *A*] and then take the limit as *A → ∞*. Since there is no terminal reproduction term and the initial state is fixed, the selection gradient becomes

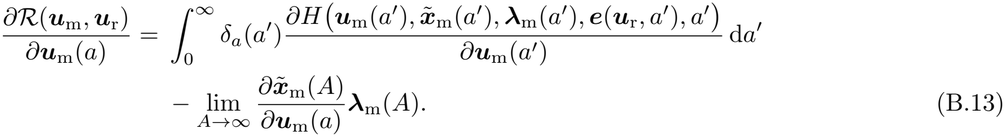

The boundary term vanishes by the transversality condition (42). Using the defining property of the Dirac delta, we therefore recover eq. (B.11) and thus eq. (B.12).

Finally, using the definition of the Hamiltonian, eq. (B.12) gives

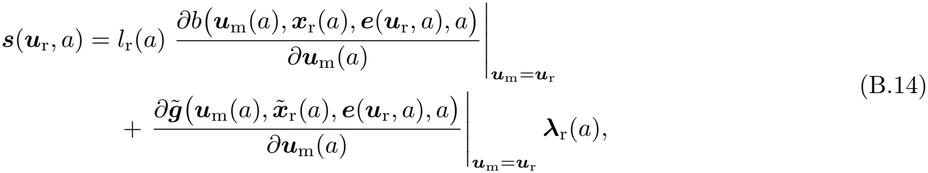

which proves eq. (44) and completes the proof of Result 5.

### B.2 Result 6

By definition, the kernel of the Jacobian operator is the functional derivative of the selection gradient with respect to the resident control,

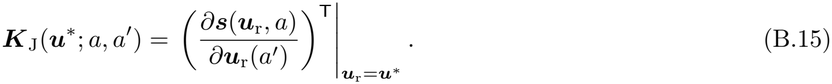

From Result 5, the selection gradient can be written as

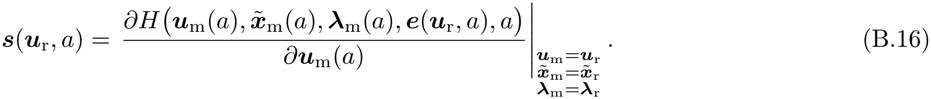

Applying the chain rule gives

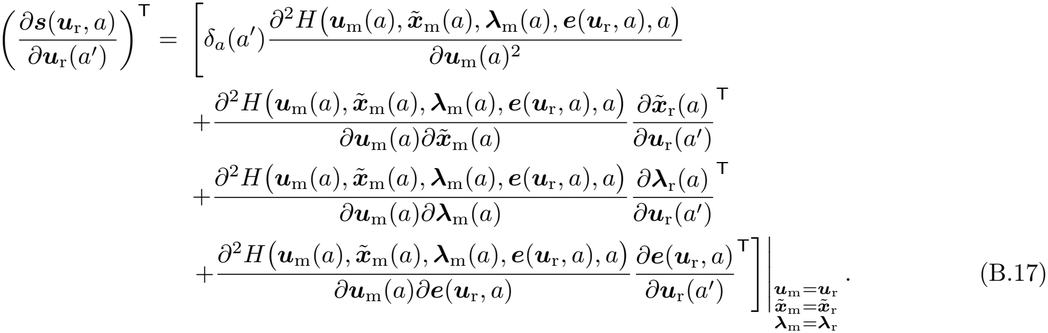

From the definition of the Hamiltonian in eq. (38),

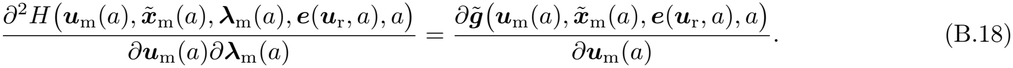

Substituting eq. (B.18) into eq. (B.17) and evaluating at the singular control, ***u***_m_ = ***u***_r_ = ***u***^*\**^, 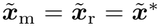, and ***λ***_m_ = ***λ***_r_ = ***λ***^*\**^, yields

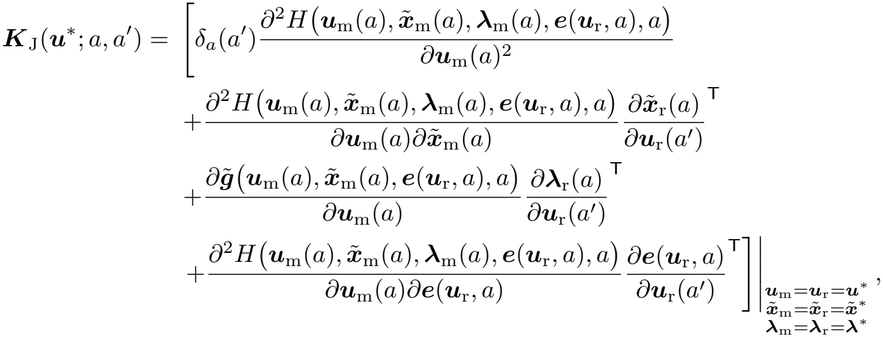

which is eq. (45) and completes the proof of Result 6.

#### B.2.1 Computing state and costate sensitivities

When explicit expressions for the resident state and costate are available as functions of the resident control, their sensitivities can be obtained directly by differentiation and substituted into the Jacobian kernel. Otherwise, these sensitivities can be computed by differentiating the resident state and costate dynamics with respect to the resident control at age *a*^*′*^. This provides a system of ODEs for 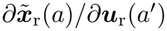and, *∂λ*_r_(*a*)/*∂u*_r_(a^*′*^) given, respectively, by

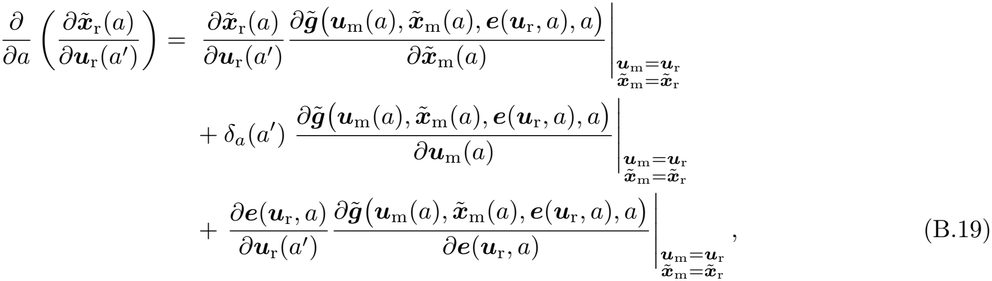

and

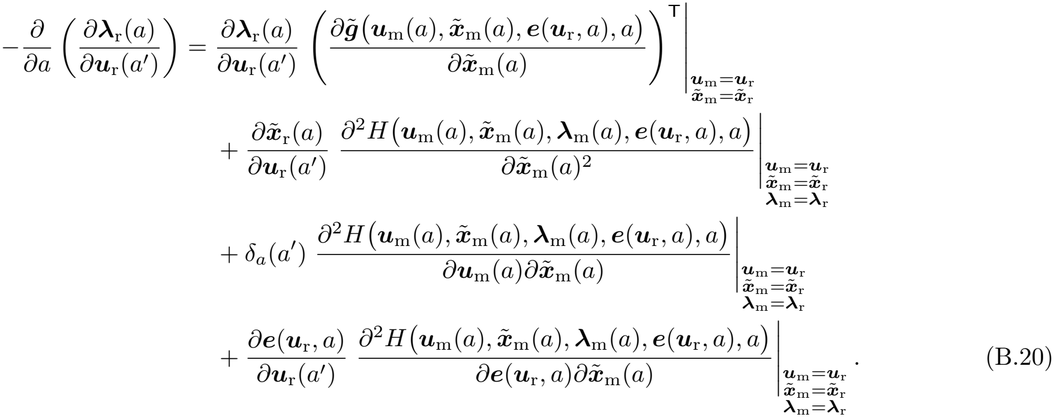

Because the initial extended state is fixed, the state sensitivity satisfies

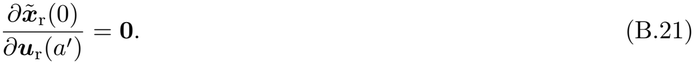

The boundary condition for the costate sensitivity is obtained by differentiating the condition defining the resident costate with respect to the resident control. For a finite lifespan, this is the terminal condition at *a*_f_, whereas for an unbounded lifespan, it follows from the appropriate asymptotic conditions, including transversality. Provided these conditions determine the costate sensitivity, and given the ecological sensitivity, the state sensitivity can be computed forward in age and the costate sensitivity backward in age.

### B.3 Result 7

By definition, the Hessian kernel is the second functional derivative of the mutant basic reproductive number with respect to the mutant control,

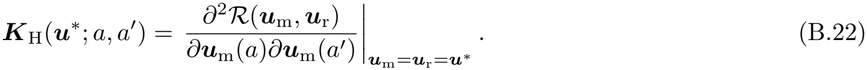

By Result 5, for both finite and unbounded lifespans

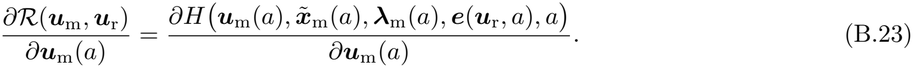

Differentiating this identity with respect to the mutant control at age *a*^*′*^, while holding the resident environment fixed, gives

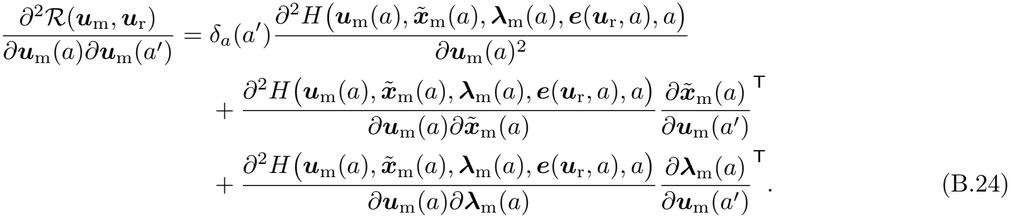

From the definition of the Hamiltonian,

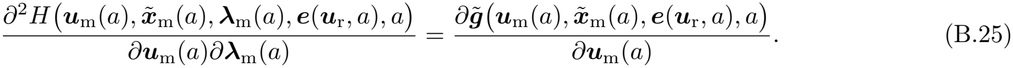

Substituting this identity into eq. (B.24) and evaluating at the singular control, ***u***_m_ = ***u***_r_ = ***u***^*\**^, 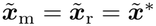, and ***λ***_m_ = ***λ***_r_ = ***λ***^*\**^, yields

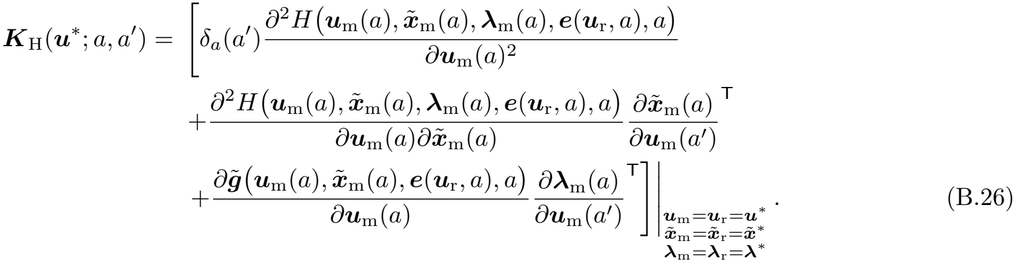

#### B.3.1 Computing state and costate sensitivities

When explicit expressions for the mutant state and costate are available as functions of the mutant control, their sensitivities can be obtained directly by differentiation and substituted into the Hessian kernel. Otherwise, these sensitivities can be computed by differentiating the mutant state and costate dynamics with respect to the mutant control at age *a*^*′*^, while holding the resident environment fixed. This then gives a system of ODEs for 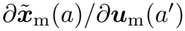 and *∂****λ***_m_(*a*)*/∂****u***_m_(*a*^*′*^), given, respectively, by

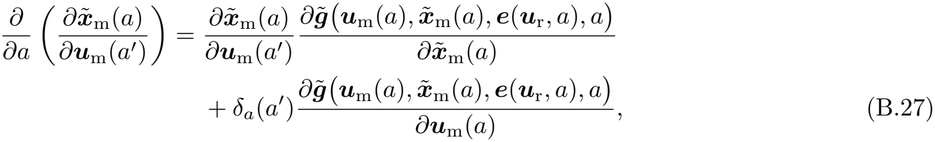

and

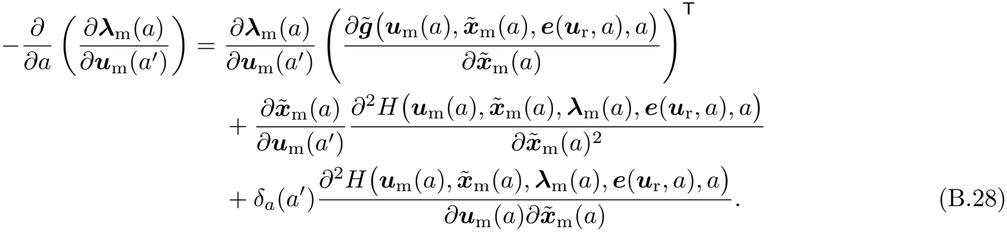

Because the initial extended state is fixed, the state sensitivity satisfies

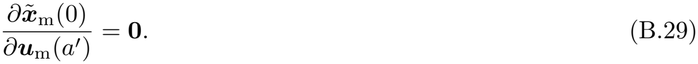

The boundary condition for the costate sensitivity is obtained by differentiating the condition defining the mutant costate with respect to the mutant control. For a finite lifespan, this is the terminal condition at *a*_f_, whereas for an unbounded lifespan, it follows from the appropriate asymptotic conditions, including transversality. Provided these conditions determine the costate sensitivity, the state sensitivity can be computed forward in age and the costate sensitivity backward in age.

#### B.3.2 Alternative representation for *K*_H_(***u***^*\**^; *a, a*^*′*^)

We here show that eq. (B.26) can be expressed only in terms of mutant state perturbations, which can be useful to know for certain applications. We only consider the finite lifespan case, *A* = [0, *a*_f_]. We start from eq. (B.2) and replace the mutant costate ***λ***_m_ by the singular costate ***λ***^*\**^, which leaves the equation valid. Substituting the resulting expression into eq. (B.22) gives

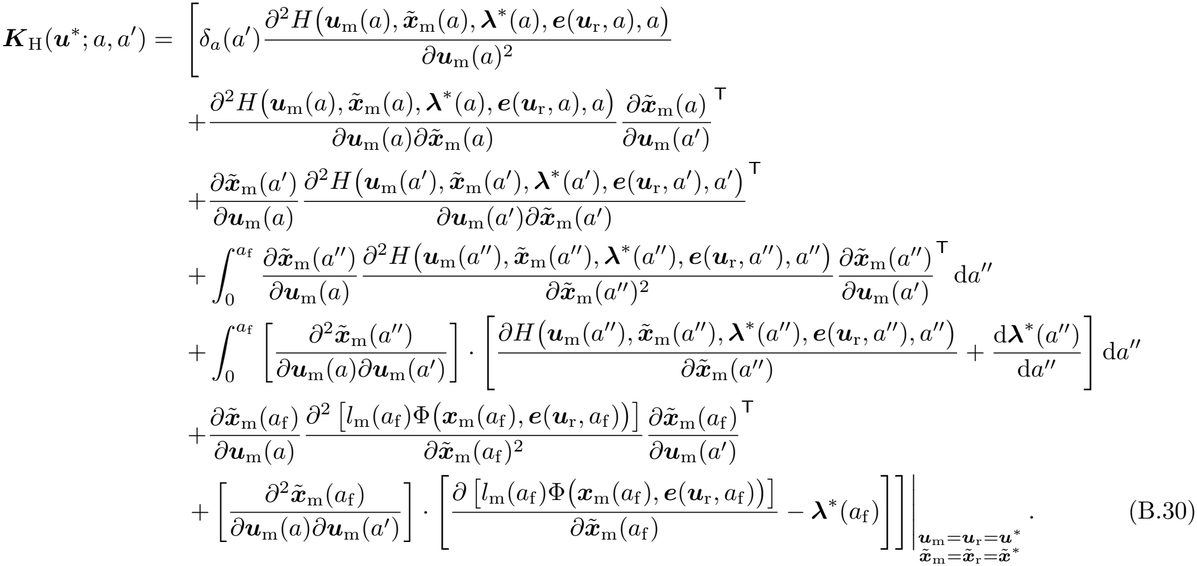

Here, 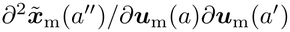 is a third-order tensor of dimension *n*_u_ *× n*_u_ *×* (1 + *n*_x_), and *·* denotes contraction over its state components. Since both terms involving these tensors vanish, we need not detail these contractions.

Indeed, the costate equation evaluated along the singular trajectory gives

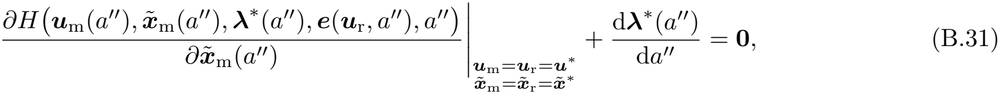

while the terminal condition gives

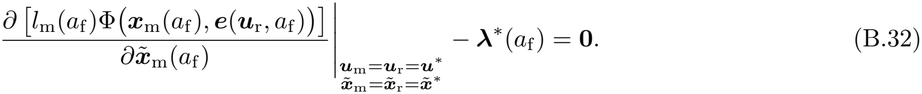

Hence,

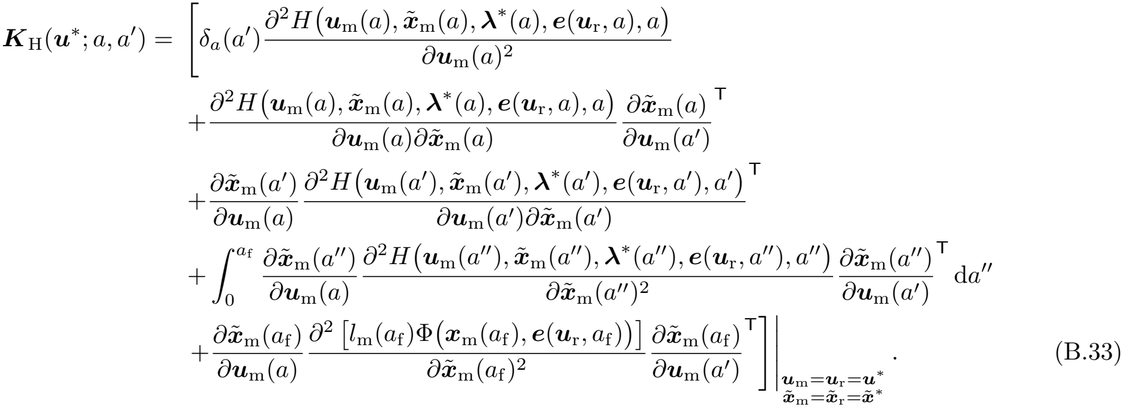

## Appendix C: Individual-based simulations

### C.1 Evolution of a floral investment schedule

Our individual-based simulations implement the flowering model introduced in section 2.6 and track the distribution of floral investment schedules in a finite population of *N* individuals over a fixed number of generations. Each individual *i* in generation *t* is characterized by its complete floral investment schedule ***u***_*i,t*_. To represent this function numerically, we divide the flowering season *A* = [0, 1] into *N*_*a*_ = 1001 equally spaced time points, *a*_*j*_ = (*j −* 1)*/*(*N*_*a*_ *−* 1), *j* ∈ *{*1, …, *N*_*a*_*}*, so that the schedule of individual *i* is represented by the vector 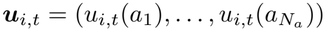.

At the first generation, all individuals express the same schedule, ***u***_*i*,1_ = 0.45*u*^*\**^ for all *i* ∈ *{*1, …, *N }*. At each generation *t ≥* 1, the following occurs.

(i) *Reproduction*. We first calculate the mean floral investment schedule expressed by the population,

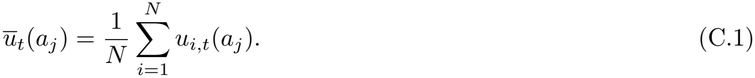

The expected fecundity of each individual is then obtained from eq. (23), with the resident schedule replaced by the current population mean schedule. Numerically, the integral over the flowering season is evaluated using the trapezoidal rule, giving the fitness of individual *i* as

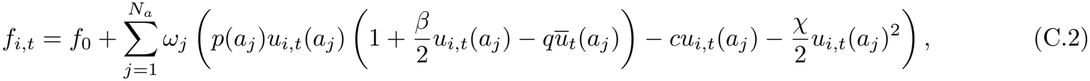

where 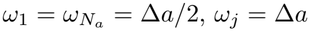 for the interior time points, and Δ*a* = 1*/*(*N*_*a*_ *−* 1). The expected fecundity *f*_*i,t*_ is then truncated at zero.

To form the next generation, we sample *N* offspring and independently assign a parent to each seed. The probability that a sampled offspring descends from individual *i* is

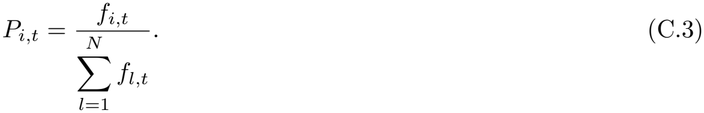

After reproduction, all adults die and are replaced by the *N* offspring, so population size remains constant.

(ii) *Mutation*. With probability 1 *− p*_mut_, an offspring inherits the complete floral investment schedule of its parent without modification. With probability *p*_mut_, its entire schedule mutates. We assume that mutational changes are smooth across the flowering season: rather than changing floral reproduction investment independently at each time point, a mutation simultaneously changes investment at all time points according to a correlated Gaussian vector.

Specifically, for each mutant offspring, we sample

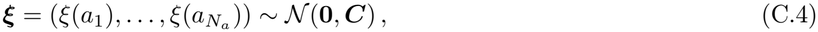

where the entries of the covariance matrix are

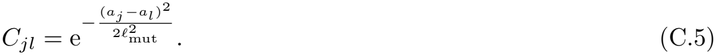

The mutant schedule is then

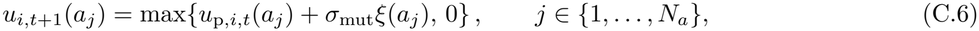

where *u*_p,*i,t*_(*a*_*j*_) is the schedule of the offspring’s parent.

Because *C*_*jj*_ = 1, the mutational effect at each time point has standard deviation *σ*_mut_. The correlation between mutational effects at two time points decreases with their separation over the flowering season, at a rate determined by *l*_mut_. Negative floral reproduction investments are truncated pointwise at the biological lower bound *u*(*a*) = 0. No upper bound is imposed.

We repeat these steps for 50,000 generations. In all simulations, we use *N* = 4000, *p*_mut_ = 0.01, *σ*_mut_ = 0.005, *l* = 0.50, *c* = 0.80, *χ* = 1.00, *f*_0_ = 2.00, and 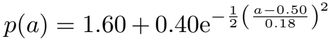. We compare a stabilizing case, with *β* = 0.35 and *q* = 0.70, to a disruptive case, with *β* = 0.80 and *q* = 1.15.

### C.2 Evolution of flower growth schedules

These simulations follow the same structure as the previous subsection, but differ in how reproductive success is calculated because it now depends on both the control and the state trajectory. At each generation, we first compute each individual’s floral-tissue trajectory 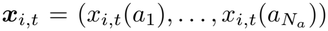 generated by its control schedule ***u***_*i,t*_, and then its fecundity. We update the state of individual *i* numerically according to

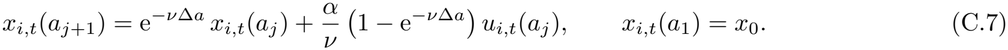

The frequency-dependent competition experienced by each individual is set by the population-mean *state*, 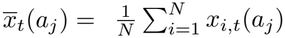. Its expected fecundity is then obtained from eq. (49) with the resident state replaced by this mean, giving, with the same trapezoidal weights *ω*_*j*_,

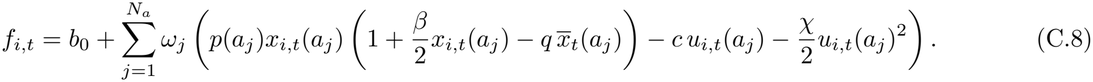

For the simulations, we used *N* = 4000, *p*_mut_ = 0.01, *σ*_mut_ = 0.005, *l*_mut_ = 0.50, *α* = 2.00, *v* = 0.70, *x*_0_ = 0, *c* = 0, *χ* = 1.00, *b*_0_ = 2.00, 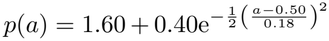. We ran the population for 100,000 generations and compared a stabilizing case, with *β* = 0.35 and *q* = 0.70, to a disruptive case, with *β* = 0.80 and *q* = 1.15.

